# Neuroticism/negative emotionality is associated with increased reactivity to uncertain threat in the bed nucleus of the stria terminalis, not the amygdala

**DOI:** 10.1101/2023.02.09.527767

**Authors:** Shannon E. Grogans, Juyoen Hur, Matthew G. Barstead, Allegra S. Anderson, Samiha Islam, Hyung Cho Kim, Manuel Kuhn, Rachael M. Tillman, Andrew S. Fox, Jason F. Smith, Kathryn A. DeYoung, Alexander J. Shackman

**Affiliations:** Departments of Psychology; Departments of Neuroscience and Cognitive Science Program; Departments of Maryland Neuroimaging Center, University of Maryland, College Park, MD 20742 USA; Department of Psychology, Yonsei University, Seoul 03722, Republic of Korea; Dead Reckoning Analytics & Consulting, College Park, MD 20740 USA; Department of Psychological Sciences, Vanderbilt University, Nashville, TN 37240 USA; Department of Psychology, University of Pennsylvania, Philadelphia, PA USA; Center for Depression, Anxiety and Stress Research, McLean Hospital, Harvard Medical School, Belmont, MA 02478 USA; Children’s National Hospital, Washington, DC 20010 USA; Department of Psychology and California National Primate Research Center, University of California, Davis, CA 95616 USA; California National Primate Research Center, University of California, Davis, CA 95616 USA

**Keywords:** neuroticism, fear and anxiety, temperament and personality, extended amygdala, bed nucleus of the stria terminalis (BST/BNST)

## Abstract

Neuroticism/Negative Emotionality (N/NE)—the tendency to experience anxiety, fear, and other negative emotions—is a fundamental dimension of temperament with profound consequences for health, wealth, and wellbeing. Elevated N/NE is associated with a panoply of adverse outcomes, from reduced socioeconomic attainment to psychiatric illness. Animal research suggests that N/NE reflects heightened reactivity to uncertain threat in the bed nucleus of the stria terminalis (BST) and central nucleus of the amygdala (Ce), but the relevance of these discoveries to humans has remained unclear. Here we used a novel combination of psychometric, psychophysiological, and neuroimaging approaches to rigorously test this hypothesis in an ethnoracially diverse, sex-balanced sample of 220 emerging adults selectively recruited to encompass a broad spectrum of N/NE. Cross-validated robust-regression analyses demonstrated that N/NE is preferentially associated with heightened BST activation during the uncertain anticipation of a genuinely distressing threat (aversive multimodal stimulation), whereas N/NE was unrelated to BST activation during certain-threat anticipation, Ce activation during either type of threat anticipation, or BST/Ce reactivity to threat-related faces. It is often assumed that different threat paradigms are interchangeable assays of individual differences in brain function, yet this has rarely been tested. Our results revealed negligible associations between BST/Ce reactivity to the anticipation of threat and the presentation of threat-related faces, indicating that the two tasks are non-fungible. These observations provide a framework for conceptualizing emotional traits and disorders; for guiding the design and interpretation of biobank and other neuroimaging studies of psychiatric risk, disease, and treatment; and for informing mechanistic research.

**SIGNIFICANCE STATEMENT:** Neuroticism/Negative Emotionality (N/NE) is a core dimension of mammalian temperament. Elevated levels of N/NE confer risk for a panoply of adversities—from reduced wealth and divorce to depression and death—yet the underlying neurobiology remains unclear. Here we show that N/NE is associated with heightened activation in the bed nucleus of the stria terminalis (BST) during the uncertain anticipation of a genuinely distressing threat. In contrast, N/NE was unrelated to BST reactivity during the certain anticipation of threat or the acute presentation of ‘threat-related’ faces, two popular probes of the emotional brain. These findings refine our understanding of what has been termed the single most important psychological risk factor in public health, with implications for on-going biobank and therapeutics research.

## INTRODUCTION

Individual differences in Neuroticism/Negative Emotionality (N/NE)—the tendency to experience anxiety and other negative emotions—have profound consequences for health, wealth, and wellbeing (<u>Shackman et al., 2016</u>). Individuals with a more negative disposition show reduced socioeconomic attainment; are more likely to experience interpersonal conflict, loneliness, unemployment, and divorce; to engage in unhealthy behaviors; to develop pathological anxiety and depression; to become physically sick; and to die prematurely (<u>Hur et al., 2019</u>; <u>Conway et al., *in press*</u>). N/NE has been conceptualized as the single most important psychological risk factor in public health, yet the underlying neurobiology remains surprisingly speculative (<u>Lahey, 2009</u>).

N/NE is thought to reflect a neurobiological tendency to overreact to threat, stressors, and other ‘trait-relevant’ challenges (<u>Shackman et al., 2016</u>; <u>Wrzus et al., 2021</u>). Although a number of circuits have been implicated, the central extended amygdala (EAc)—including the central nucleus of the amygdala (Ce) and bed nucleus of the stria terminalis (BST)—has received the most empirical scrutiny and occupies a privileged position in most theoretical models (<u>Fox et al., 2015a</u>; <u>DeYoung et al., 2022</u>; <u>Kagan, 2022</u>). Mechanistic work demonstrates that the EAc is critical for orchestrating defensive responses to a variety of threats in humans and other animals (<u>Fox and Shackman, 2019</u>; <u>Zhu et al., 2024</u>). Neuroimaging studies in monkeys show that Ce and BST reactivity to uncertain threat covaries with trait-like variation in anxious temperament and behavioral inhibition, core facets of N/NE (<u>Fox et al., 2015b</u>; <u>Shackman et al., 2017</u>). But the relevance of these discoveries to humans remains unclear. Only a handful of human studies have used genuinely distressing threats to assess relations between N/NE and EAc function and most have focused exclusively on the amygdala proper (**Table 1**). Even less is known about the BST. Only one small-scale study has directly addressed this question, providing preliminary evidence that individuals with a more negative disposition show heightened BST activation during the uncertain anticipation of aversive stimulation (<u>Somerville et al., 2010</u>). Although modest samples and a lack of attention to the BST preclude decisive inferences, these observations motivate the hypothesis that N/NE reflects heightened recruitment of the BST, and possibly the Ce, during the anticipation of aversive stimulation and suggest that these associations may be more pronounced when threat is uncertain.

**Table 1.** Human studies of N/NE and distress-eliciting neuroimaging paradigms. ^a^Older normalization techniques (e.g., affine) can introduce substantial spatial smoothing and registration error, which is a concern for work focused on small subcortical structures, such as the Ce and BST. Abbreviations—AFNI, Analysis of Functional NeuroImages software package; BBR, boundary-based registration of the T1- and T2-weighted images; BFI-N, Big Five Inventory-Neuroticism; EAc, central extended amygdala; fROI, functionally defined region-of-interest (all others are anatomically defined); NEO-FFI, Neuroticism/Extraversion/Openness Five-Factor Inventory-3; NR, not reported or indeterminate; NS, not significant; SPM#, Statistical Parametric Mapping software package (# indicates version); STAI, Spielberger State-Trait Anxiety Inventory; SVC, Small-Volume Corrected; TT, piecewise linear Talairach-Tournoux transformation; UCS, Unconditioned Stimulus.

| Study | N (% Male) | Neuroticism/Negative Emotionality (N/NE) | MRI Field Strength (T) | EPI Voxel Size (mm<sup>3</sup>) | Normalization<sup>a</sup> | Trait-Relevant Challenge(s) | Analysis Type | Summary of EAc Results |
| --- | --- | --- | --- | --- | --- | --- | --- | --- |
| **Present Study** | 220 (49.5%), stratified sampling to capture a broad spectrum of N/NE; screened to exclude current internalizing illness | Multi-Scale, Multi-Occasion Composite | 3 | 10.5 | BBR and Diffeomorphic | Anticipation of Uncertain/Certain Aversive Shock, Auditory Clips, and Photographs;<br>Negative Emotional Faces vs. Place Scenes | ROIs, Voxelwise | See the main report |
| **Brinkmann et al., 2018** (Brinkmann et al., 2018) | 93 (35.5%), screened to exclude any internalizing illness in the past five years | STAI | 3 | 15.3 | Affine and Manual TT | Aversive Photographs | NR | **NS** |
| **Indovina et al., 2011** (Indovina et al., 2011) | 23 (43.5%), screened to exclude lifetime internalizing illness | STAI | 3 | 18.0 | "SPM5" | Pavlovian Threat Cue (Auditory UCS; Uninstructed) | Amygdala ROI | **Positive relations** between the magnitude of amygdala reactivity and N/NE |
| **Kirlic et al., 2019** (Kirlic et al., 2019) | 83 (42.2%), 43 of whom had depression or anxiety disorder diagnoses | Multi-Scale Composite | 3 | 18.1 | "AFNI" and Diffeomorphic | Anticipation of Uncertain Shock | Voxelwise | **NS** |
| **Klumpers et al., 2017** (Klumpers et al., 2017) | 108 (100%), screened to exclude lifetime internalizing illness | STAI | 1.5 | 38.1 | "SPM8" | Anticipation of Uncertain Shock | Amygdala and BST ROIs | **NS** |
| **Schuyler et al., 2012** (Schuyler et al., 2012) | 127 (36.2%), screened to exclude past-year internalizing illness | BFI-N | 3 | 56.25 | Affine | Aversive Photographs | Amygdala fROI | **Positive relations** between the persistence of amygdala reactivity and N/NE |
| **Sjouwerman et al., 2020** (Sjouwerman et al., 2020) | 113 (61.1%), screened to exclude lifetime internalizing illness | STAI | 3 | 8.0 | "SPM8" | Pavlovian Threat Cue (Shock UCS; Uninstructed) | Amygdala SVC | **Positive relations** between amygdala reactivity and N/NE |
| **Somerville et al., 2010** (Somerville et al., 2010) | 50 (44%), screened to exclude lifetime internalizing illness | Multi-Scale Composite | 3 | 31.5 | "SPM2" | Anticipation of Uncertain Shock | Amygdala and BST fROIs, Voxelwise | **Positive relations** between the magnitude of BST reactivity and N/NE<br>**NS Amygdala** |
| **Visser et al., 2021** (Visser et al., 2021) | 98 (30.6%), screened to exclude lifetime internalizing illness | STAI | 3 | 8.0 | BBR and Diffeomorphic | Pavlovian Threat Cue (Shock UCS; Uninstructed) | Amygdala SVC | **NS Amygdala** |

Here we used fMRI to quantify EAc reactivity to a genuinely distressing threat-anticipation paradigm and test its relevance to N/NE in an ethnoracially diverse sample of 220 emerging adults (**Figures 1-2**). Participants were selectively recruited from a pool of 6,594 pre-screened individuals, ensuring a broad spectrum of N/NE. Prior neuroimaging studies of N/NE have relied on overly optimistic analytic approaches (<u>Marek et al., 2022</u>). Here we used anatomically defined Ce and BST regions-of-interest (ROIs) and cross-validated robust estimates of brain-temperament associations (**Figure 1**). To enhance power, we leveraged a composite measure of N/NE—aggregated across two scales and three measurement occasions—to minimize fluctuations in responding (<u>Nikolaidis et al., 2022</u>; <u>Roemer et al., *in press*</u>) (**Figure 3**). Hypotheses and approach were pre-registered.

**Figure 1.**
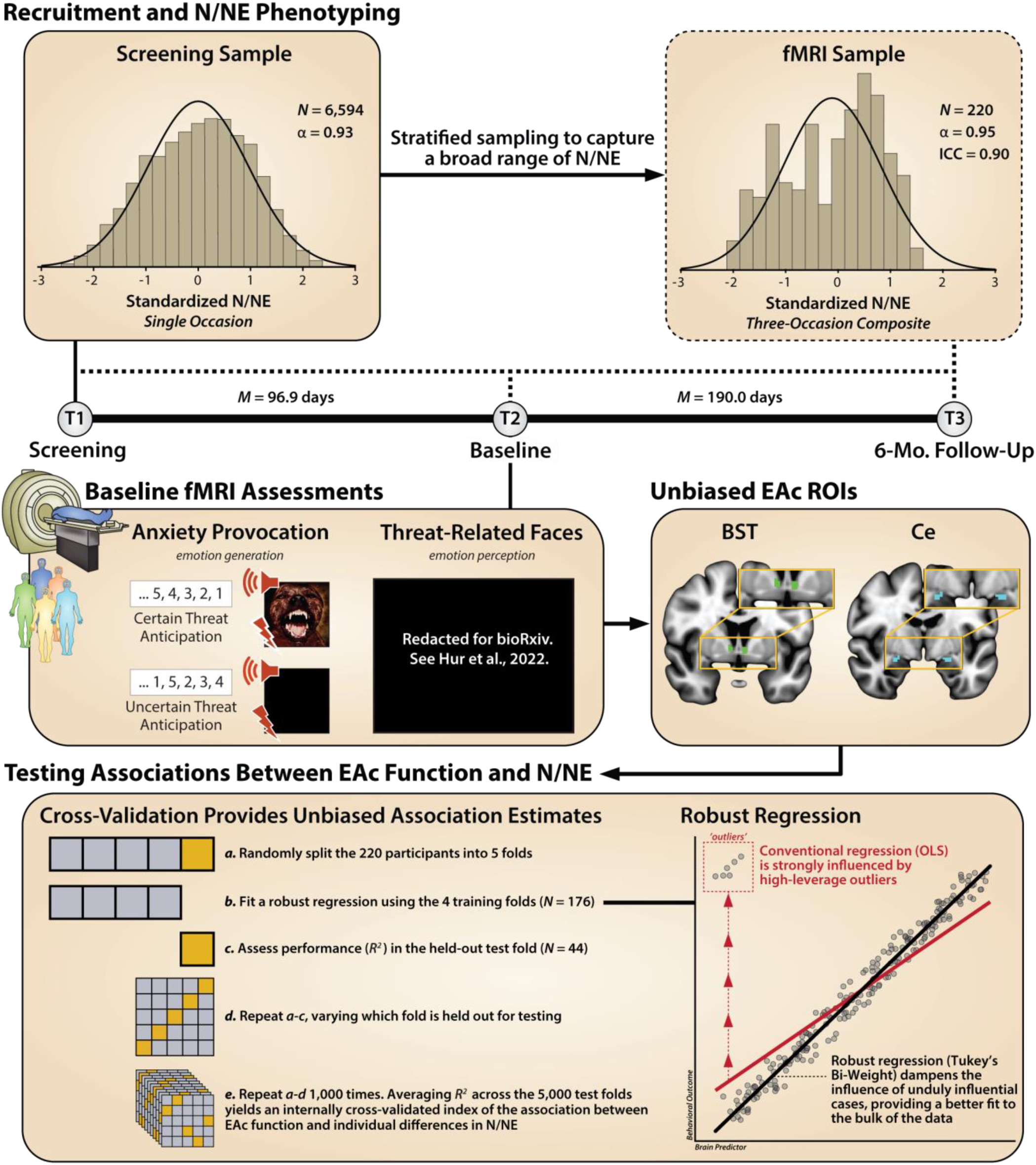
Study overview. ***<u>Recruitment and N/NE Phenotyping.</u>*** To ensure a broad spectrum of N/NE, participants were selectively recruited from an ethnoracially diverse pool of 6,594 pre-screened individuals. N/NE was assessed at screening (T1), at the baseline laboratory session (T2), and at the 6-month follow-up session (T3). To maximize reliability and power, analyses leveraged a composite measure of N/NE that was aggregated across 2 scales and 3 measurement occasions. Top panels indicate the distribution (histogram), internal-consistency reliability (α), and test-retest reliability of N/NE in the screening (*left*) and fMRI (*right*) samples. Timeline indicates the interval between assessments. ***<u>Baseline fMRI Assessments.</u> Anxiety Provocation.*** Following the baseline laboratory assessment (T2), participants completed an fMRI assessment. All participants completed the Maryland Threat Countdown paradigm, a well-established anxiety-provocation paradigm. The paradigm takes the form of a 2 (*Valence:* Threat/Safety) × 2 (*Temporal Certainty:* Certain/Uncertain) factorial design. On threat trials, subjects saw a stream of integers that terminated with the temporally certain or uncertain presentation of a noxious electric shock, unpleasant photograph, and thematically related audio clip. Safety trials were similar but terminated with the delivery of benign stimuli. Hypothesis testing focused on neural activation associated with the anticipation of temporally certain and uncertain threat, relative to safety. A total of 220 individuals provided usable imaging data. ***Threat-Related Faces.*** A subset of 213 participants also completed a ‘threat-related’ (fearful/angry) faces fMRI paradigm. Participants viewed short blocks of photographs, alternating between blocks of faces and places (e.g., park, office). Hypothesis testing focused on activation associated with threat-related faces, relative to places. ***<u>EAc ROIs.</u>*** Anatomically defined regions-of-interest (ROIs) enabled us to rigorously test the central hypothesis that N/NE reflects heightened recruitment of the BST (*green*), and potentially the Ce (dorsal amygdala; *cyan*), during aversive the anticipation of a genuinely aversive threat, and explore the possibility that these associations are more evident when the timing of threat encounters is uncertain. Unlike conventional whole-brain voxelwise analyses—which screen thousands of voxels for statistical significance and yield optimistically biased associations—anatomically defined ROIs ‘fix’ the measurements-of-interest *a priori*, providing statistically unbiased estimates of brain-phenotype associations. Standardized regression coefficients were extracted and averaged across voxels for each combination of ROI, task contrast, and participant. ***<u>Testing Associations Between EAc Function and N/NE.</u> Cross-Validation Provides Statistically Unbiased Association Estimates***. Conventional-regression approaches use all available data for model fitting (‘training’), yielding optimistically biased estimates of model performance (*R^2^*) that do not generalize well to unseen data (‘overfitting’). As shown in the bottom-left panel, we used a well-established cross-validation framework (i.e., repeated 5-fold) to compute statistically unbiased associations. ***Robust Regression.*** As shown in the bottom-right panel, conventional regression is sensitive to high-leverage outliers (*red*). Here we used robust regression (Tukey’s bi-weight) to reduce the influence of unduly influential cases, providing a better fit to the bulk of the data, and reducing volatility across the cross-validated training (*N*=176) and test (*N*=44) folds. The same analytic framework was used for the faces paradigm. Abbreviations—α, Cronbach’s alpha (internal-consistency reliability); BST, bed nucleus of the stria terminalis; Ce, dorsal amygdala in the region of the central nucleus; EAc, central extended amygdala; ICC, intraclass correlation (test-retest reliability); *M*, mean; Mo., months; *N*, number of observations; N/NE, neuroticism/negative emotionality; OLS, ordinary least squares; ROI, region of interest.

**Figure 2.**
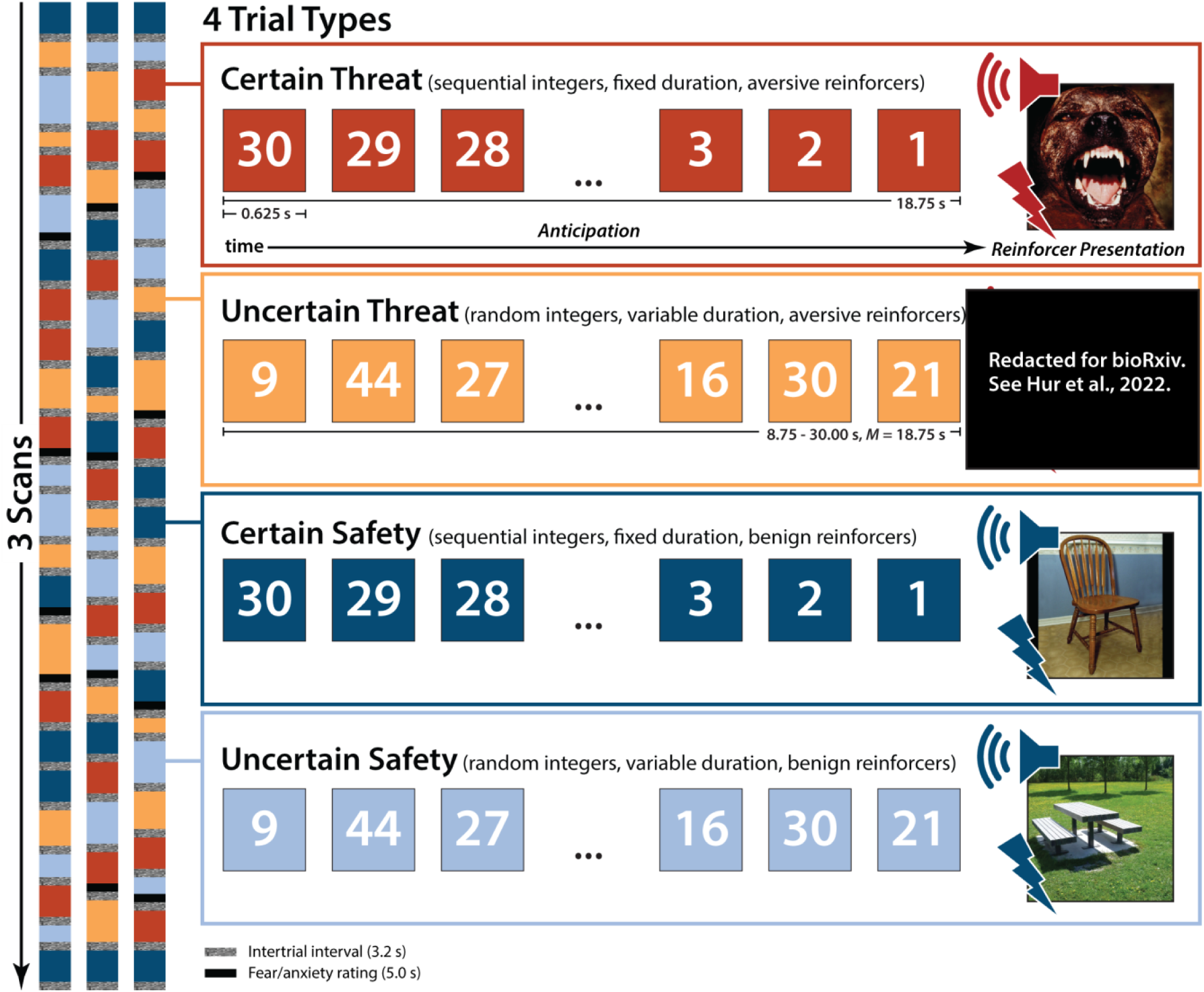
Threat-Anticipation paradigm. The Maryland Threat Countdown paradigm takes the form of a 2 (*Valence:* Threat/Safety) × 2 (*Temporal Certainty:* Certain/Uncertain) repeated-measures, randomized event-related design. Participants were completely informed about the task design and contingencies prior to scanning. The task was administered in 3 scans, with short breaks between scans. On certain-threat trials, participants saw a descending stream of integers (‘count-down’) for 18.75 s. To ensure robust anxiety, this anticipation epoch always terminated with the presentation of a noxious electric shock, unpleasant photograph, and thematically related audio clip (e.g., scream). Uncertain-threat trials were similar, but the integer stream was randomized and presented for an uncertain and variable duration (8.75-30.00 s; *M*=18.75 s). Participants knew that something aversive was going to occur, but they had no way of knowing precisely *when*. Safety trials were similar, but terminated with the delivery of benign reinforcers (e.g., just-perceptible electrical stimulation). White-noise visual masks (3.2 s) were presented between trials to minimize persistence of visual reinforcers in iconic memory. Participants were periodically prompted to rate the intensity of fear/anxiety experienced a few seconds earlier, during the anticipation (‘countdown’) epoch of the prior trial. Skin conductance was continuously acquired.

**Figure 3.**
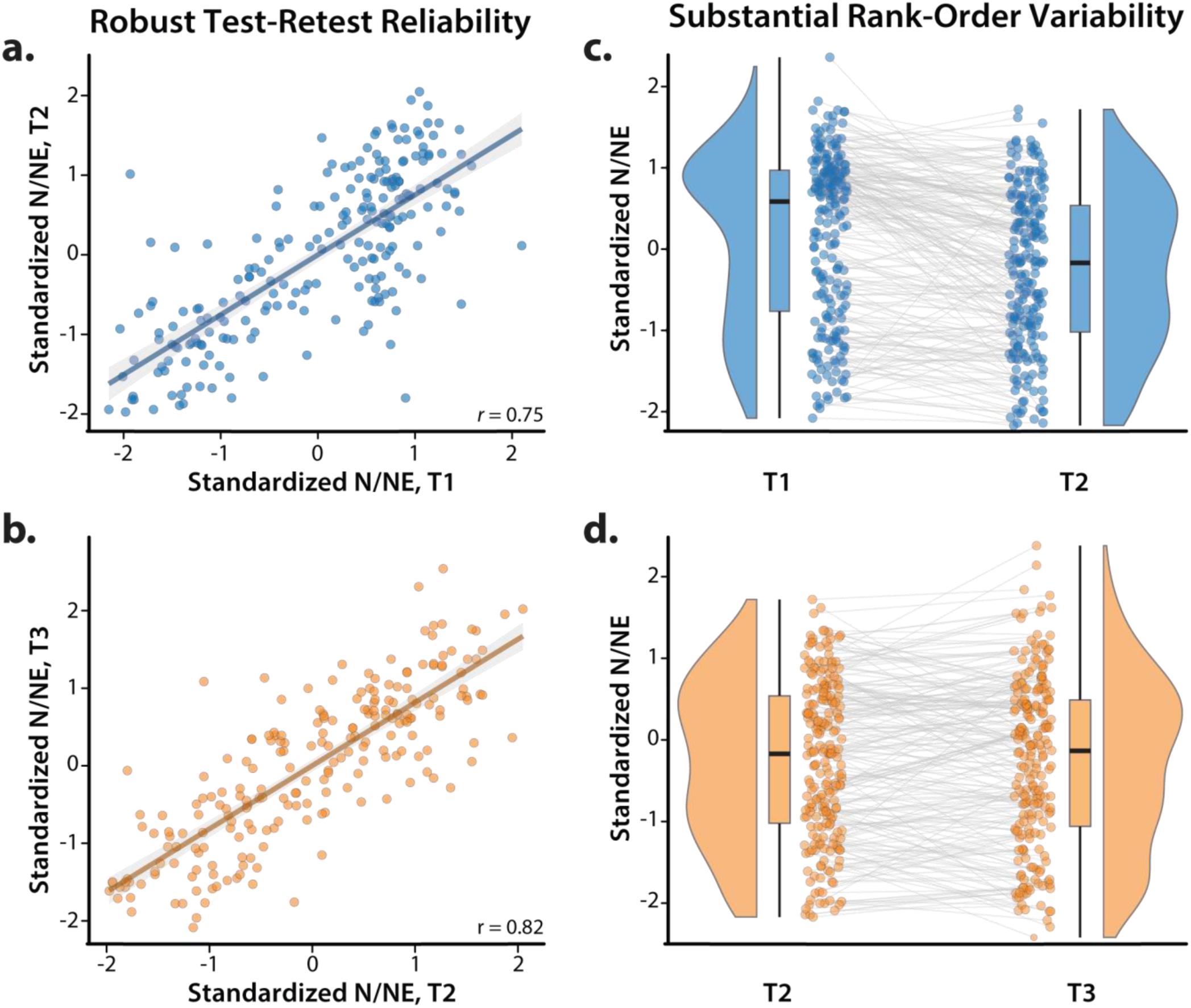
Robust test-retest reliability belies substantial rank-order variability in N/NE. A voluminous literature underscores the reliability of questionnaire measures of personality and temperament (<u>Soto and John, 2017</u>). Indeed, as shown in panels *a* and *b*, our multiscale measure of N/NE showed robust test-retest reliability across successive measurement occasions (cf. Figure 1). Nevertheless, inspection of the raincloud plots depicted in panels *c* and *d* revealed notable variability in the rank order of participants, with 46.4% of the sample showing at least a 0.5 *SD* change between T1 (Screening) and T2 (Baseline), 41.4% between T2 and T3 (6-Month Follow-Up), and 55.9% between T1 and T3. In short, even highly reliable single-occasion measurements of N/NE contain considerable state and error variance (<u>Roemer et al., *in*</u> *<u>press</u>*). Consistent with recent methodological recommendations (<u>Nikolaidis et al., 2022</u>), we addressed this issue by creating a composite measure of N/NE that was aggregated across 2 scales and 3 measurement occasions (T1-T3), maximizing reliability and power to detect brain-trait associations. Scatterplots depict standard (ordinary least squares) regression estimates. Gray bands indicate 95% confidence intervals. Raincloud plots depict the smoothed density distributions (i.e., ‘bean’ or ‘half-violin’) of standardized N/NE. Box-and-whisker plots indicate the medians (*horizontal lines*) and interquartile ranges (*boxes*). Whiskers depict 1.5× the interquartile range. Colored dots connected by gray lines indicate changes in standardized N/NE for each participant.

To provide a more direct link with on-going research, we performed parallel analyses using data from an overlapping sample that completed an emotional-faces paradigm (**Figure 4**). Variants of this paradigm are widely used in biobank research, often in the guise of assessing Research Domain Criteria (RDoC) ‘Negative Valence Systems’ (<u>Tozzi et al., 2020</u>; <u>Grogans et al., 2022</u>). Although photographs of ‘threat-related’ facial expressions robustly activate the EAc, they do not elicit substantial distress and are better conceptualized as an index of threat perception (<u>Hur et al., 2019</u>). The inclusion of the two threat paradigms also allowed us to test whether they are statistically interchangeable. It is often tacitly assumed that different tasks targeting a common function (e.g., ‘threat’) are more-or-less equivalent probes of individual differences in brain function. Yet this assumption of convergent validity has rarely been examined empirically, never in a large sample, and never in the BST (<u>Villalta-Gil et al., 2017</u>).

**Figure 4.**
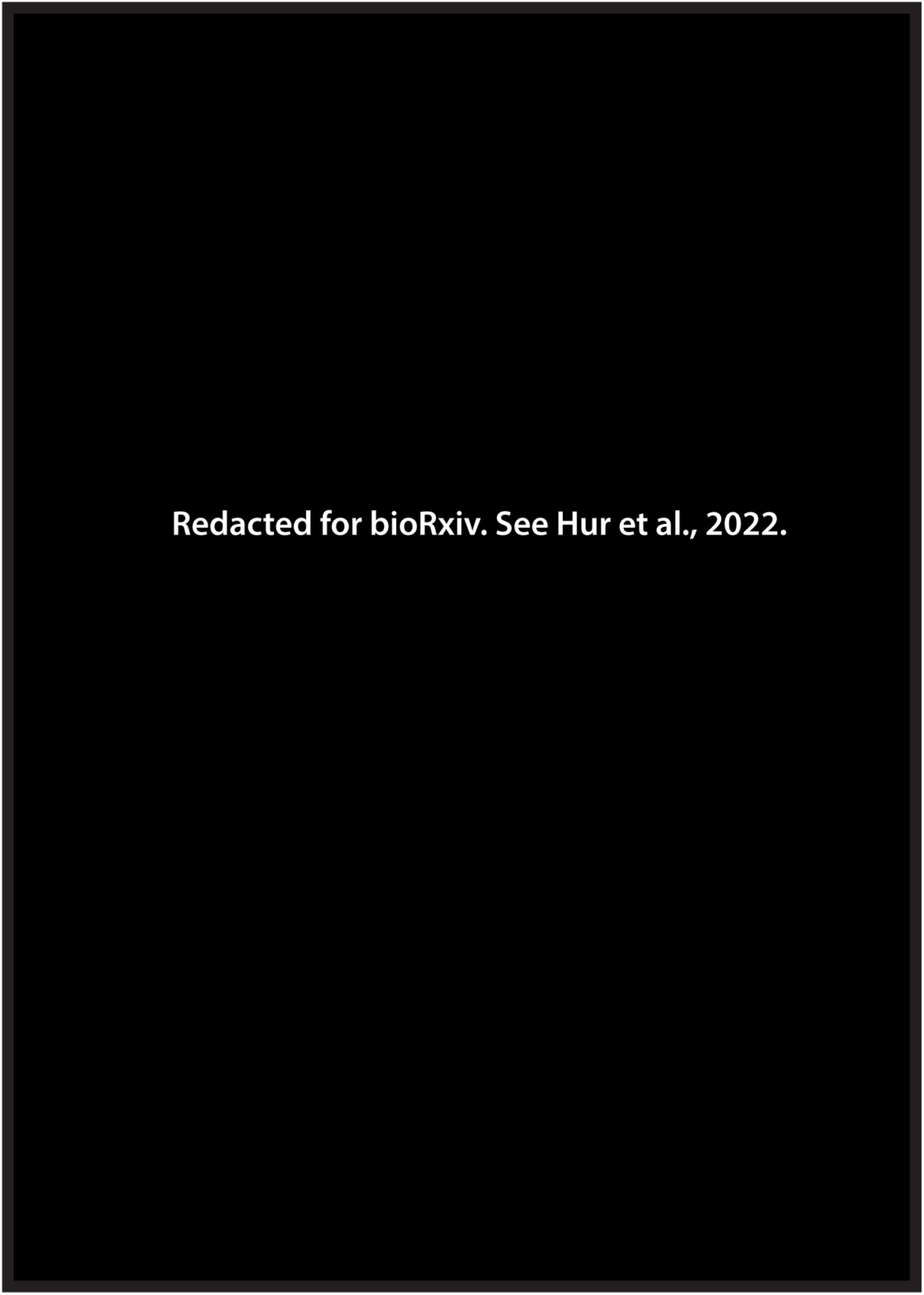
Threat-perception (*emotional-faces) paradigm.* The emotional-faces paradigm took the form of a pseudo-randomized block design and was administered in 2 scans. During each scan, participants viewed standardized photographs of adults (half female) modeling prototypical angry faces, fearful faces, happy faces, or places (i.e., emotionally neutral everyday scenes; 7 blocks/condition/scan). Blocks consisted of 10 briefly presented photographs of faces or places (1.6 s) separated by fixation crosses (0.4 s). To minimize potential habituation, each photograph was presented a maximum of two times. To ensure engagement, on each trial, participants judged whether the current photograph matched that presented on the prior trial (i.e., performed a ‘1-back’ continuous-performance task).

## METHOD

### Overview

As part of a recently completed prospective-longitudinal study focused on individuals at risk for the development of internalizing disorders (R01-MH107444), we used well-established psychometric measures of N/NE to screen 6,594 first-year university students (57.1% female; 59.0% White, 19.0% Asian, 9.9% African American, 6.3% Hispanic, 5.8% Multiracial/Other; *M*=19.2 years, *SD*=1.1 years) (<u>Shackman et al., 2018</u>). Screening data were stratified into quartiles (top quartile, middle quartiles, bottom quartile), separately for males and females. Individuals who met preliminary inclusion criteria were independently and randomly recruited via email from each of the resulting six strata. Given our focus on psychiatric risk, approximately half the participants were recruited from the top quartile, with the remainder split between the middle and bottom quartiles (i.e., 50% high, 25% medium, and 25% low). This enabled us to sample a broad spectrum of N/NE without gaps or discontinuities—in contrast to prior work focused on convenience samples (<u>Charpentier et al., 2021</u>)—while balancing the inclusion of men and women (cf. **Figure 1**). Simulation work suggests that this over-sampling (‘enrichment’) approach does not bias statistical tests to a degree that would compromise their validity (<u>Hauner et al., 2014</u>). All participants had normal or corrected-to-normal color vision, and reported the absence of lifetime neurological symptoms, pervasive developmental disorder, very premature birth, medical conditions that would contraindicate MRI, and prior experience with noxious electrical stimulation. All participants were free from a lifetime history of psychotic and bipolar disorders; a current diagnosis of a mood, anxiety, or trauma disorder (past 2 months); severe substance abuse; active suicidality; and on-going psychiatric treatment as determined by an experienced, masters-level diagnostician using the Structured Clinical Interview for DSM-5 (<u>First et al., 2015</u>). Participants provided informed written consent and all procedures were approved by the Institutional Review Board at the University of Maryland, College Park (Protocol #659385). Data from this study were featured in prior work focused on relations between personality and the longitudinal course of internalizing symptoms (<u>Conway et al., *in press*</u>), validation of the threat-anticipation paradigm (<u>Hur et al., 2020b</u>; <u>Shackman et al., *accepted in principle*</u>), relations between social anxiety and real-world mood (<u>Huret al., 2020a</u>), relations between threat-related brain activity and real-world mood dynamics (<u>Hur et al., 2022</u>), and the neuroanatomical correlates of early-life anxiety, shyness, and behavioral inhibition (<u>Bas-Hoogendam et al., 2022</u>), but have never been used to address the present aims.

### Power Analyses

Sample size was determined *a priori* as part of the application for the grant that supported data collection (R01-MH107444). The target sample size (*N*≈240) was chosen to afford acceptable power and precision given available resources. At the time of study design, G-power (version 3.1.9.2) indicated >99% power to detect a benchmark (‘generic’) medium-sized effect (*r*=0.30) with up to 20% planned attrition (*N*=192 usable datasets) using α=0.05, two-tailed (<u>Erdfelder et al., 1996</u>). The final sample of 220 usable fMRI datasets (see below) provides 80% power to detect a focal brain-temperament association as small as *r*=0.186 (*R^2^*=3.46%).

### Participants

A total of 241 participants were recruited and scanned. Of these, 6 withdrew due to excess distress in the scanner, 1 withdrew from the study after the imaging session, and 4 were excluded due to incidental neurological findings.

#### Threat-anticipation paradigm

One participant was excluded from fMRI analyses due to gross susceptibility artifacts in the echoplanar imaging (EPI) data, 2 were excluded due to insufficient usable data (see below), 6 were excluded due to excess motion artifact (see below), and 1 was excluded due to task timing issues, yielding a racially diverse final sample of 220 participants (49.5% female; 61.4% White, 18.2% Asian, 8.6% African American, 4.1% Hispanic, 7.3% Multiracial/Other; *M*=18.8 years, *SD*=0.4). Of these, 2 participants were excluded from skin conductance analyses due to insufficient usable data (see below).

#### Threat-perception (emotional-faces) paradigm

Three participants were excluded due to gross susceptibility artifacts in the EPI data, 1 was excluded due to insufficient usable data (see below), 7 were excluded due to excessive motion artifact (see below), and 6 participants for inadequate behavioral performance (see below), yielding a final sample of 213 participants (49.3% female; 61.0% White, 17.8% Asian, 8.5% African American, 4.2% Hispanic, 7.0% Multiracial/Other; *M*=18.8 years, *SD*=0.3). A subset of 209 participants (98.1%) provided usable data for both fMRI tasks and were used for the cross-task association analyses.

### Neuroticism/Negative Emotionality (N/NE)

#### Broadband N/NE

As in our prior work (<u>Shackman et al., 2018</u>; <u>Hur et al., 2020a</u>; <u>Hur et al., 2020b</u>; <u>Hur et al., 2022</u>), we used 2 well-established measures of neuroticism (Big Five Inventory-Neuroticism) (<u>John et al., 2008</u>) and trait anxiety (International Personality Item Pool-Trait Anxiety) (<u>Goldberg, 1999</u>; <u>Goldberg et al., 2006</u>) to quantify individual differences in N/NE on three occasions: screening, baseline, and 6-month follow-up (see **Figure 1** in the main report). Participants used a 1 (*disagree strongly*) to 5 (*agree strongly*) scale to rate a total of 18 items (e.g., *depressed or blue, tense, worry, nervous, get distressed easily, fear for the worst, afraid of many things*). At screening, the neuroticism and anxiety scales were strongly correlated (*r*s>0.85) and internally consistent (*α*s>0.85). To minimize the influence of short-term fluctuations in responding, hypothesis testing employed a multi-scale, multi-occasion composite measure of N/NE. This was computed by z-transforming the neuroticism and trait-anxiety scales (using the mean and variance from the much larger screening sample) and then averaging across scales and measurement occasions. The resulting composite captured a sizable range of the N/NE spectrum and demonstrated strong internal-consistency (*α*=0.95) and test-retest (ICC*_3,k_*=0.90) reliability (cf. **Figures 1** and **3).**

#### N/NE Facets

Epidemiological, psychiatric, and biological studies typically focus on broadband measures of N/NE, including the Eysenck Personality Questionnaire; Spielberger’s State-Trait Anxiety Inventory; and our own multiscale composite (see above) (<u>Shackman et al., 2016</u>; <u>Hur et al., 2019</u>). Yet it is clear that N/NE is a complex phenotype that subsumes several narrower traits, including dispositional fear/anxiety, depression/sadness, and anger/irritability (<u>Caspi et al., 2005</u>; <u>Soto and John, 2017</u>). N/NE is also associated with elevated emotional volatility (<u>Soto and John, 2017</u>; <u>Kalokerinos et al., 2020</u>). Basic affective neuroscience research suggests that each of these facets of N/NE reflects partially distinct neural circuits, and recent research demonstrates a mixture of shared and unique psychological associations and biological substrates (<u>Soto and John, 2017</u>; <u>Fox et al., 2018</u>; <u>Thorp et al., 2021</u>; <u>Klein-Flügge et al., 2022</u>; <u>Watson et al., 2022</u>; <u>Khoo et al., 2023</u>). To understand which facets of N/NE are most closely associated with EAc reactivity, we leveraged the revised Big Five Inventory (BFI-2), a well-established, hierarchically organized scale that was purposely constructed to enable psychometrically rigorous facet-level analyses (<u>Soto and John, 2017</u>). At the baseline and 6-month follow-up sessions (but not the screening assessment), participants used a 1 (*disagree strongly*) to 5 (*agree strongly*) scale to rate 12 items (4 items/facet) indexing dispositional Anxiety (e.g., *anxious or afraid*), Depression/Sadness (e.g., *sad*), and Emotional Volatility (e.g., *easily upset, moody*). Paralleling the approach used for broadband N/NE, facet scores were averaged across assessments to minimize occasion-specific fluctuations. All three facets demonstrated adequate internal-consistency (α=0.81-0.88) and test-retest (*r*=0.69-0.79) reliability. As expected, all were robustly correlated with our broadband composite (*r*=0.74-0.92). Inter-facet associations, while substantial, indicated substantial unique variance (*r*=0.68-0.71; *R^2^*<51%).

### Threat-Anticipation Paradigm

#### Paradigm Structure and Design Considerations

The Maryland Threat Countdown paradigm is a well-established, fMRI-optimized variant of temporally uncertain-threat assays that have been validated using fear-potentiated startle and acute anxiolytic administration (e.g., benzodiazepine) in mice, rats, and humans (<u>Miles et al., 2011</u>; <u>Hefner et al., 2013</u>; <u>Daldrup et al., 2015</u>; <u>Lange et al., 2017</u>; <u>Moberg et al., 2017</u>). The paradigm has been successfully deployed in a sample of university students that overlaps the sample featured here (<u>Hur et al., 2020b</u>; <u>Hur et al., 2022</u>) and in an independent sample of community volunteers (<u>Kim et al., 2023</u>).

As shown in **Figure 2**, the Maryland Threat Countdown takes the form of a 2 (*Valence:* Threat/Safety) × 2 (*Temporal Certainty:* Uncertain/Certain) randomized, event-related, repeated-measures design (3 scans; 6 trials/condition/scan). Subjects were completely informed about the task design and contingencies prior to scanning. Simulations were used to optimize the detection and deconvolution of task-related hemodynamic signals. Stimulus presentation and ratings acquisition were controlled using Presentation software (version 19.0, Neurobehavioral Systems, Berkeley, CA).

On certain-threat trials, participants saw a descending stream of integers (‘count-down;’ e.g., 30, 29, 28…3, 2, 1) for 18.75 s. To ensure robust distress, this anticipation epoch culminated with the presentation of a noxious electric shock, unpleasant photograph (e.g., mutilated body), and thematically related audio clip (e.g., scream, gunshot). Uncertain-threat trials were similar, but the integer stream was randomized and presented for an uncertain and variable duration (8.75-30.00 s; *M*=18.75 s). Participants knew that something aversive was going to occur, but they had no way of knowing precisely when. Consistent with methodological recommendations (<u>Shackman and Fox, 2016</u>), the average duration of the anticipation epoch was identical across conditions, ensuring an equal number of measurements (TRs/condition). The specific mean duration was chosen to enhance detection of task-related differences in the blood oxygen level-dependent (BOLD) signal (‘activation’) (<u>Henson, 2007</u>) and to allow sufficient time for sustained responses to become evident. Safety trials were similar, but terminated with the delivery of benign reinforcers (see below). Valence was continuously signaled during the anticipation epoch (‘countdown’) by the background color of the display. Temporal certainty was signaled by the nature of the integer stream. Certain trials always began with the presentation of the number 30. On uncertain trials, integers were randomly drawn from a near-uniform distribution ranging from 1 to 45 to reinforce the impression that they could be much shorter or longer than certain trials and to minimize incidental temporal learning (‘time-keeping’). To concretely demonstrate the variable duration of uncertain trials, during scanning, the first three trials featured short (8.75 s), medium (15.00 s), and long (28.75 s) anticipation epochs. To mitigate potential confusion and eliminate mnemonic demands, a lower-case ‘c’ or ‘u’ was presented at the lower edge of the display throughout the anticipatory epoch. White-noise visual masks (3.2 s) were presented between trials to minimize the persistence of visual reinforcers in iconic memory.

Participants were periodically prompted (following the offset of the white-noise visual mask) to rate the intensity of fear/anxiety experienced a few seconds earlier, during the anticipation (‘countdown’) period of the prior trial, using a 1 (*minimal*) to 4 (*maximal*) scale and an MRI-compatible response pad (MRA, Washington, PA). Each condition was rated once per scan (16.7% trials). Premature ratings (<300 ms) were censored. All participants provided at least 6 usable ratings and rated each condition at least once. Skin conductance was continuously acquired throughout.

#### Procedures

Prior to scanning, participants practiced an abbreviated version of the paradigm (without electrical stimulation) until they indicated and staff confirmed understanding. Benign and aversive electrical stimulation levels were individually titrated. *Benign Stimulation.* Participants were asked whether they could “reliably detect” a 20 V stimulus and whether it was “at all unpleasant.” If the subject could not detect the stimulus, the voltage was increased by 4 V and the process repeated. If the subject indicated that the stimulus was unpleasant, the voltage was reduced by 4 V and the process was repeated. The final level chosen served as the benign electrical stimulation during the imaging assessment (*M*=21.06 V, *SD*=5.06). *Aversive Stimulation.* Participants received a 100 V stimulus and were asked whether it was “as unpleasant as you are willing to tolerate”—an instruction specifically chosen to maximize anxious distress and arousal. If the subject indicated that they were willing to tolerate more intense stimulation, the voltage was increased by 10 V and the process repeated. If the subject indicated that the stimulus was too intense, the voltage was reduced by 5 V and the process repeated. The final level chosen served as the aversive electrical stimulation during the imaging assessment (*M*=117.85 V, *SD*=26.10). The intensity of aversive stimulation was weakly and negatively associated with individual differences in N/NE (*r*(218)=-0.13, *p*=0.05). Following each scan, staff re-assessed whether stimulation was sufficiently intense and increased the level as necessary.

#### Electrical Stimuli

Electrical stimuli (100 ms; 2 ms pulses every 10 ms) were generated using an MRI-compatible constant-voltage stimulator system (STMEPM-MRI; Biopac Systems, Inc., Goleta, CA). Stimuli were delivered using MRI-compatible, disposable carbon electrodes (Biopac) attached to the fourth and fifth digits of the non-dominant hand.

#### Visual Stimuli

A total of 72 aversive and benign photographs (1.8 s) were selected from the International Affective Picture System (<u>for details, see Hur et al., 2020b</u>). Visual stimuli were digitally back-projected (Powerlite Pro G5550, Epson America, Inc., Long Beach, CA) onto a semi-opaque screen mounted at the head-end of the scanner bore and viewed using a mirror mounted on the head-coil.

#### Auditory Stimuli

A total of 72 aversive and benign auditory stimuli (0.8 s) were adapted from open-access online sources. Auditory stimuli were delivered using an amplifier (PA-1 Whirlwind) with in-line noise-reducing filters and ear buds (S14; Sensimetrics, Gloucester, MA) fitted with noise-reducing ear plugs (Hearing Components, Inc., St. Paul, MN).

#### Skin Conductance

Skin conductance was continuously acquired during each scan using a Biopac system (MP-150; Biopac Systems, Inc., Goleta, CA). Skin conductance (250 Hz; 0.05 Hz high-pass) was measured using MRI-compatible disposable electrodes (EL507) attached to the second and third digits of the non-dominant hand.

### Threat-Perception (Emotional-Faces) Paradigm

The emotional-faces paradigm took the form of a pseudo-randomized block design and was administered in 2 scans, with a short break between scans (**Figure 4**). During each scan, participants viewed standardized photographs of adults (half female) modeling prototypical angry faces, fearful faces, happy faces, or places (i.e., emotionally neutral everyday scenes; 7 blocks/condition/scan). To maximize signal strength and homogeneity and mitigate potential habituation (<u>Henson, 2007</u>; <u>Maus et al., 2010</u>; <u>Plichta et al., 2014a</u>; <u>Plichta et al., 2014b</u>), each 20-s block consisted of 10 briefly presented photographs of faces or places (1.6 s) separated by fixation crosses (0.4 s). To minimize potential habituation, each photograph was presented a maximum of two times (<u>for details, see Hur et al., 2020b</u>). To ensure engagement, on each trial, participants judged whether the current photograph matched that presented on the prior trial (i.e., a ‘1-back’ continuous-performance task). Matches occurred 37.1% of the time.

### MRI Data Acquisition

MRI data were acquired using a Siemens Magnetom TIM Trio 3 Tesla scanner (32-channel head-coil). During scanning, foam inserts were used to immobilize the participant’s head within the head-coil and mitigate potential motion artifact. Participants were continuously monitored using an MRI-compatible eye-tracker (Eyelink 1000; SR Research, Ottawa, Ontario, Canada) and the AFNI real-time motion plugin (<u>Cox, 1996</u>). Sagittal T1-weighted anatomical images were acquired using a magnetization prepared rapid acquisition gradient echo sequence (TR=2,400 ms; TE=2.01 ms; inversion time=1,060 ms; flip=8°; slice thickness=0.8 mm; in-plane=0.8 × 0.8 mm; matrix=300 × 320; field-of-view=240 × 256). A T2-weighted image was collected co-planar to the T1-weighted image (TR=3,200 ms; TE=564 ms; flip angle=120°). To enhance resolution, a multi-band sequence was used to collect oblique-axial EPI volumes (multiband acceleration=6; TR=1,250 ms; TE=39.4 ms; flip=36.4°; slice thickness=2.2 mm, number of slices=60; in-plane resolution=2.1875 × 2.1875 mm; matrix=96 × 96). Images were collected in the oblique-axial plane (approximately −20° relative to the AC-PC plane) to minimize potential susceptibility artifacts. For the threat-anticipation task, three 478-volume EPI scans were acquired. For the threat-perception (emotional-faces) task, two 454-volume EPI scans were acquired. The scanner automatically discarded 7 volumes prior to the first recorded volume. To enable fieldmap correction, two oblique-axial spin echo (SE) images were collected in opposing phase-encoding directions (rostral-to-caudal and caudal-to-rostral) at the same location and resolution as the functional volumes (i.e., co-planar; TR=7,220 ms; TE=73 ms). Measures of respiration and pulse were continuously acquired during scanning using a respiration belt and photo-plethysmograph affixed to the first digit of the non-dominant hand. Following the last scan, participants were removed from the scanner, debriefed, compensated, and discharged.

### Skin Conductance Data Processing Pipeline

Skin conductance data were processed using *PsPM* (version 4.0.2) and in-house Matlab (version 9.9.0.1467703) code (<u>Bach and Friston, 2013</u>; <u>Bach et al., 2018</u>). Data were orthogonalized with respect to pulse and respiration signals and de-spiked using *filloutliers* (150-sample moving-median widow; modified Akima cubic Hermite interpolation). Each scan was then band-pass filtered (0.009-0.333 Hz), median centered, and down-sampled (4 Hz). Subject-specific skin conductance response functions (SCRFs) were estimated by fitting the four parameters of the canonical SCRF (<u>Bach et al., 2010</u>) to the grand-average reinforcer response using *fmincon* and a cost function that maximized variance explained and penalized negative coefficients.

### MRI Data Processing Pipeline

Methods were optimized to minimize spatial normalization error and other potential sources of noise. Data were visually inspected before and after processing for quality assurance.

#### Anatomical Data Processing

Methods are similar to those described in other recent reports by our group (<u>Hur et al., 2020b</u>; <u>Hur et al., 2022</u>; <u>Kim et al., 2023</u>). T1-weighted images were inhomogeneity corrected using *N4* (<u>Tustison et al., 2010</u>) and denoised using *ANTS* (<u>Avants et al., 2011</u>). The brain was then extracted using a combination of *BEaST* (<u>Eskildsen et al., 2012</u>) and brain-extracted-and-normalized reference-brains (<u>BIAC, 2022</u>). Brain-extracted T1 images were normalized to a version of the brain-extracted 1-mm T1-weighted MNI152 (version 6) template (<u>Grabner et al., 2006</u>) modified to remove extracerebral tissue. Normalization was performed using the diffeomorphic approach implemented in *SyN* (version 2.3.4) (<u>Avants et al., 2011</u>). T2-weighted images were rigidly co-registered with the corresponding T1 prior to normalization. The brain-extraction mask from the T1 was then applied. Tissue priors were unwarped to native space using the inverse of the diffeomorphic transformation (<u>Lorio et al., 2016</u>). Brain-extracted T1 and T2 images were segmented—using native-space priors generated in *FAST* (version 6.0.4) (<u>Jenkinson et al., 2012</u>)—for subsequent use in T1-EPI co-registration (see below).

#### Fieldmap Data Processing

SE images and *topup* were used to create fieldmaps. Fieldmaps were converted to radians, median-filtered, and smoothed (2-mm). The average of the distortion-corrected SE images was inhomogeneity corrected using *N4* and masked to remove extracerebral voxels using *3dSkullStrip* (version 19.1.00).

#### Functional Data Processing

EPI files were de-spiked using *3dDespike*, slice-time corrected to the TR center using *3dTshift*, and motion corrected to the first volume and inhomogeneity corrected using *ANTS* (12-parameter affine). Transformations were saved in ITK-compatible format for subsequent use (<u>McCormick et al., 2014</u>). The first volume was extracted for EPI-T1 co-registration. The reference EPI volume was simultaneously co-registered with the corresponding T1-weighted image in native space and corrected for geometric distortions using boundary-based registration (<u>Jenkinson et al., 2012</u>). This step incorporated the previously created fieldmap, undistorted SE, T1, white matter (WM) image, and masks. To minimize incidental spatial blurring, the spatial transformations necessary to transform each EPI volume from native space to the reference EPI, from the reference EPI to the T1, and from the T1 to the template were concatenated and applied to the processed EPI data in a single step. Normalized EPI data were resampled (2 mm^3^) using fifth-order b-splines. Hypothesis testing focused on anatomically defined EAc regions of interest (ROIs), as detailed below. To maximize anatomical resolution, no additional spatial filters were applied, consistent with prior work by our team and recent recommendations (<u>Tillman et al., 2018</u>; <u>Kim et al., 2023</u>).

### Skin Conductance Data Exclusions and Modeling

#### Data Exclusions

To ensure data validity, ‘scans’ (i.e., runs) that did not show numerically positive skin conductance responses to reinforcer (e.g., shock) presentation (averaged across trials) were censored. Participants with <2 usable scans were excluded from skin conductance analyses (see above).

#### First-Level Modeling

Robust general linear models (GLMs) were used to separate electrodermal signals associated with the threat anticipation (‘countdown’) epochs from those evoked by other aspects of the task (e.g., reinforcer delivery). Modeling was performed separately for each participant and scan using *robustfit*. Subject-specific SCRFs were convolved with rectangular regressors time-locked to the presentation of the reinforcers (separately for each trial type), visual masks, and rating prompts. To quantify skin conductance level (SCL) during the anticipation epochs, first-level residuals were averaged separately for each participant and condition.

### fMRI Data Exclusions, Modeling, and Derivative Variables

#### Data Exclusions

Volume-to-volume displacement (>0.5 mm) was used to assess residual motion artifact. Scans with excessively frequent artifacts (>2 *SD*) were discarded. Participants with insufficient usable fMRI data (<2 scans of the threat-anticipation task or <1 scan of the threat-perception task) or who showed poor behavioral performance on the threat-perception task (see above; accuracy <2 *SD*) were excluded from the relevant analyses (see above).

#### First-Level (Single-Subject) fMRI Modeling

For each participant, first-level modeling was performed using GLMs implemented in *SPM12* (version 7771), with the default autoregressive model and the temporal band-pass filter set to the hemodynamic response function (HRF) and 128 s (<u>Wellcome Centre for Human Neuroimaging, 2022</u>). Regressors were convolved with a canonical HRF. *Threat-Anticipation Paradigm.* Hemodynamic reactivity was modeled using variable-duration rectangular (‘boxcar’) regressors that spanned the anticipation (‘countdown’) epochs of uncertain-threat, certain-threat, and uncertain-safety trials. To maximize design efficiency, certain-safety anticipation served as the reference condition and contributed to the baseline estimate (<u>Poline et al., 2007</u>). Epochs corresponding to the presentation of the four types of reinforcers, white-noise visual masks, and rating prompts were simultaneously modeled using the same approach. EPI volumes acquired before the first trial and following the final trial were unmodeled and contributed to the baseline estimate. Consistent with prior work (<u>Hur et al., 2020b</u>; <u>Hur et al., 2022</u>; <u>Kim et al., 2023</u>), nuisance variates included estimates of volume-to-volume displacement, motion (6 parameters × 3 lags), cerebrospinal fluid (CSF) signal, instantaneous pulse and respiration rates, and ICA-derived nuisance signals (e.g., brain edge, CSF edge, global motion, white matter) (<u>Pruim et al., 2015</u>). Volumes with excessive volume-to-volume displacement (>0.5 mm) and those during and immediately following reinforcer delivery were censored. *Threat-Perception (Emotional-Faces) Paradigm.* Hemodynamic reactivity to blocks of each emotional expression (angry, fearful, and happy) was modeled using time-locked rectangular regressors. Place blocks served as the reference condition and contributed to the baseline estimate (<u>Poline et al., 2007</u>).

#### EAc ROIs

Consistent with prior work by our group, task-related Ce and BST activation was quantified using well-established, anatomically defined ROIs and spatially unsmoothed fMRI data (<u>Tillman et al., 2018</u>; <u>Kim et al., 2023</u>) (cf. **Figure 1**). The derivation of the Ce ROI is detailed in (<u>Tillman et al., 2018</u>). The probabilistic BST ROI was developed by Theiss and colleagues and thresholded at 25% (<u>Theiss et al., 2017</u>). It mostly encompasses the supra-commissural BST, given the difficulty of reliably discriminating the borders of regions below the anterior commissure in T1-weighted images (<u>Kruger et al., 2015</u>). Bilateral ROIs were decimated to the 2-mm resolution of the fMRI data. EAc ROI analyses used standardized regression coefficients extracted and averaged for each contrast (e.g., Uncertain-Threat anticipation), region, and participant. Unlike conventional whole-brain voxelwise analyses—which screen thousands of voxels for statistical significance and yield optimistically biased associations—anatomically defined ROIs ‘fix’ the measurements-of-interest *a priori*, providing statistically unbiased estimates of brain-phenotype associations (<u>Poldrack et al., 2017</u>).

### Analytic Strategy

#### Overview

Analyses were performed using a combination of *SPM12* (<u>Wellcome Centre for Human Neuroimaging, 2022</u>), *R* (version 4.0.2), *Rstudio* (version 1.2.1335)*, JASP* (version 0.17.1), and *<u>jamovi</u>* (version 2.3.16) (<u>Love et al., 2019</u>; <u>R Core Team, 2022</u>; <u>RStudio Team, 2022</u>; <u>The jamovi project, 2023</u>). Some analyses were performed using *psych* (version 2.2.5) (<u>Revelle, 2022</u>) and *stats* (version 4_4.0.2) (<u>R Core Team, 2022</u>). Diagnostic procedures and data visualizations were used to confirm that test assumptions were satisfied (<u>Tukey, 1977</u>). Some figures were created using *ggplot2* (version 3.3.6) (<u>Wickham, 2016</u>), *raincloudplots* (version 0.2.0) (<u>Allen et al., 2021</u>), and *MRIcron* (<u>Rorden, 2019</u>). Clusters and peaks were labeled using the Harvard–Oxford atlas (<u>Frazier et al., 2005</u>; <u>Desikan et al., 2006</u>; <u>Makris et al., 2006</u>).

#### Confirmatory Testing

As a precursor to hypothesis testing, we used standard repeated-measures GLMs to confirm that the threat-anticipation paradigm amplified subjective symptoms of distress (in-scanner fear/anxiety ratings) and objective signs of arousal (SCL). These analyses also afforded an opportunity to explore whether individual differences in N/NE amplify reactivity to a well-controlled, genuinely distressing experimental challenge (Maryland Threat Countdown) (<u>Shackman et al., 2016</u>). Using the multiscale, multi-occasion composite (see above), mean-centered N/NE was included as a fully crossed dimensional covariate. Significant interactions were decomposed using focal contrasts and regressions. Exploratory nonparametric tests yielded similar conclusions (not reported).

We also confirmed that the threat-anticipation paradigm recruited the EAc. To maximize reliability, we focused on estimates of activation (e.g., uncertain-threat anticipation) relative to the implicit baseline (certain safety anticipation) (<u>Brown et al., 2011</u>; <u>Infantolino et al., 2018</u>; <u>Baranger et al., 2021</u>; <u>Heilicher et al., 2022</u>; <u>Kennedy et al., 2022</u>). Spatially unsmoothed data and whole-brain voxelwise (‘second-level’) repeated-measures GLMs (‘random effects’) were used to separately examine certain and uncertain threat (vs. implicit baseline). We also examined the broader contrast of threat relative to safety anticipation. Significance was assessed using FDR *q*<.05 (whole-brain corrected). A series of one-sample Student’s *t*-tests was used to confirm that the anatomically defined EAc ROIs (BST and Ce) showed nominally significant activation during certain- and uncertain-threat anticipation (*p*<0.05, uncorrected). A standard 2 (*Region:* BST, Ce) × 2 (*Certainty:* Certain, Uncertain) repeated-measures GLM was used to explore potential regional differences in sensitivity to certain-versus-uncertain threat anticipation. The same general approach was used to confirm that threat-related faces (vs. implicit baseline) recruit the EAc (BST and Ce). Whole-brain statistical parametric maps are freely available at Neurovault.org (https://neurovault.org/collections/13109). Processed ROI data are freely available at the Open Science Framework (https://osf.io/w5cdk).

#### Hypothesis Testing and Exploratory Analyses

The primary aim of the present study was to test the hypothesis that variation in N/NE is associated with heightened recruitment of the BST, and potentially the Ce, during the anticipation of threat, and determine whether this association is more evident when the timing of threat encounters is uncertain. Two robust regression models were implemented for each ROI (BST and Ce). The first quantified relations between individual differences in N/NE—indexed using the multi-scale, multi-occasion composite—and general threat reactivity, independent of the temporal certainty of aversive stimulation (threat vs. safety anticipation). The second model quantified relations between N/NE and uncertain-threat reactivity, controlling for certain-threat reactivity. This provides an estimate of the variance in N/NE uniquely explained by BST or Ce activation during the anticipation of temporally uncertain (and/or certain) threat.

Conventional-regression approaches use all available data for model testing, yielding optimistically biased estimates of performance (*R^2^*) that do not generalize well to unseen data (model ‘overfitting’) (<u>Marek et al., 2022</u>; <u>Spisak et al., 2023</u>). To address this, we used *k*-folds cross-validation (5-folds × 1,000 repetitions) to compute statistically unbiased estimates of brain-temperament associations (cross-validated *R^2^*) (<u>Poldrack et al., 2020</u>). Conventional regression is also sensitive to stochastic noise (e.g., sampling and measurement error). This can be conceptualized as another form of model overfitting. Estimates derived from such models can change dramatically given relatively small changes in the data, rendering them intrinsically less reproducible (<u>Yarkoni and Westfall, 2017</u>; <u>Yu and Yao, 2017</u>). Here we instead used a robust-regression approach (Tukey’s bi-weight), which reduces the impact of unduly influential cases, provides a better fit to the bulk of the data, and reduces volatility across the cross-validated training (*N*=176) and test (*N*=44) folds (**Figure 1**) (<u>Yu and Yao, 2017</u>). Robust- and conventional-regression approaches show similar performance when conventional-model assumptions are met (<u>Kafadar, 1983</u>; <u>Wager et al., 2005</u>; <u>Yu and Yao, 2017</u>).

Hypothesis testing was implemented using *caret* (version 6.0-86) and the default Tukey biweight loss function (*c*=4.685), which provides 95% asymptotic efficiency when conventional-regression assumptions are satisfied (<u>Kafadar, 1983</u>; <u>Kuhn et al., 2020</u>). By convention, final models were trained using all available data. To clarify specificity, we used the same analytic framework to quantify relations between N/NE and EAc reactivity to threat-related faces. Conventional-regression results are available at the Open Science Framework (https://osf.io/w5cdk). Sensitivity analyses confirmed that our key conclusions remained unchanged when controlling for potential nuisance variation in the mean-centered intensity of self-selected aversive electrical stimulation (not reported).

When significant associations between EAc threat reactivity and N/NE were detected, follow-up analyses were used to compare the strength of association across brain metrics (<u>Hotelling, 1940</u>) and explore potential associations with individual differences in the intensity of task-elicited distress (mean-centered fear/anxiety ratings). We also explored the relevance of narrower emotional traits, indexed using the mean-centered BFI-2 Anxiety, Depression/Sadness, and Emotional Volatility facet scales (averaged across the baseline and 6-month follow-up assessments). Separate robust GLMs were computed for each facet scale.

Whole-brain voxelwise GLMs (random effects) were used to explore associations between mean-centered N/NE and activation in less intensely scrutinized regions. No significant voxelwise associations were detected (FDR *q*<0.05, corrected).

The second aim of the present study was to determine whether threat-anticipation and threat-perception are interchangeable probes of individual differences in EAc function, that is, whether the two fMRI paradigms show ‘convergent validity’ (<u>Villalta-Gil et al., 2017</u>). Using the same analytic framework detailed above, we computed robust between-task correlations, separately for the BST and Ce ROIs, for each of the key imaging contrasts. Demonstrating that individual differences in BST and/or Ce activation during the anticipation of threat (aversive multimodal stimulation) are strongly correlated with individual differences in activation during the presentation of threat-related (angry/fearful) faces would provide evidence of cross-task convergence.

### Preregistration and Deviation

Our general approach and hypotheses were preregistered (<u>Grogans and Shackman, 2021</u>). The preregistration called for a conventional regression approach. To enhance rigor, we adopted a cross-validated robust-regression framework for hypothesis testing. Recent work underscores the necessity of cross-validation for reproducible estimates of brain-phenotype associations (<u>Spisak et al., 2023</u>).

## RESULTS

### Threat anticipation amplifies subjective distress and objective arousal

We used standard repeated-measures GLMs to confirm that the threat-anticipation paradigm had the intended impact on anxious distress and arousal. Mean-centered N/NE was included as a fully crossed dimensional covariate, allowing us to explore the possibility that variation in N/NE influences reactivity to this well-controlled emotional challenge (<u>Shackman et al., 2016</u>).

As shown in **Figure 5a**, fearful and anxious feelings were significantly elevated during the anticipation of threat compared to safety, and this was particularly evident when the timing of aversive encounters was uncertain (*Valence: F*(1,218)=1,135.06, *p*<0.001; *Certainty: F*(1,218)=212.95, *p*<0.001; *Valence × Certainty: F*(1,218)=31.75, *p*<0.001; *Threat, Uncertain vs. Certain: F*(1,218)=148.90, *p*<0.001; *Safety, Uncertain vs. Certain: F*(1,218)=78.78, *p*<0.001).

**Figure 5.**
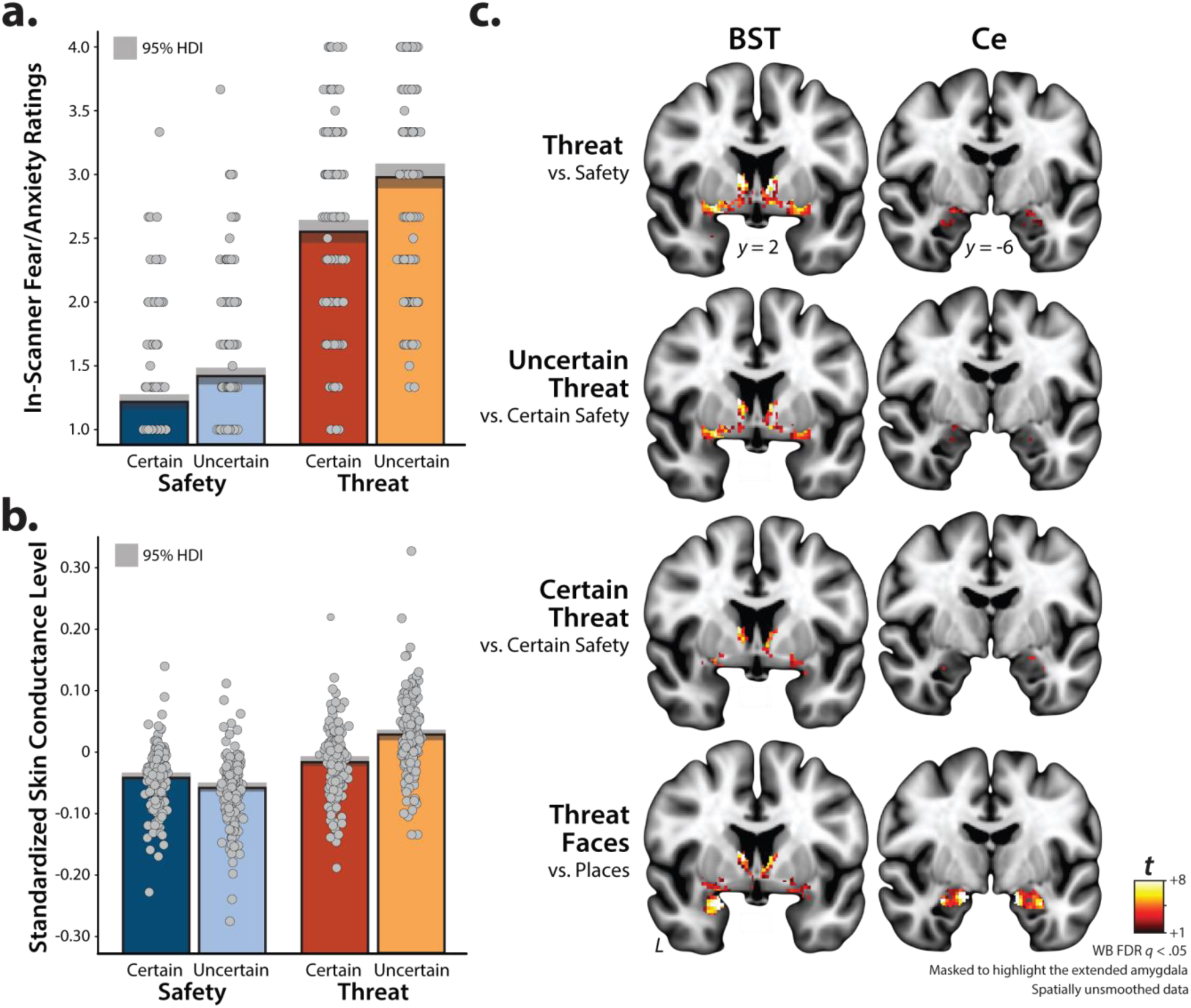
The threat-anticipation and -perception tasks are valid probes of EAc function. **a. *<u>Threat anticipation evokes subjective distress.</u>*** Fear and anxiety were increased during the anticipation of threat compared to safety, and this was particularly evident for temporally uncertain threat (*p*<0.001). **b. *<u>Threat anticipation evokes objective signs of arousal.</u>*** A similar pattern was evident for skin conductance level (*p*<0.001). **c. *<u>Threat-anticipation and threat-perception recruit the EAc.</u>*** As shown in the top three rows, the anticipation of threat, uncertain threat, and certain threat recruited the BST and the dorsal amygdala in the region of the Ce, when compared to their respective reference conditions (*q*<0.05, corrected). As shown in the bottom row, the acute presentation of threat-related faces was also associated with significant activation in the region of the BST and the dorsal amygdala (Ce), relative to the reference condition. For additional details, see **Supplemental Tables 5-1, 5-2, 5-3,** and **5-4** (https://osf.io/w5cdk). Bars indicate the means (*colored bars*), Bayesian 95% highest density intervals (*gray bands*), and individual participants (*gray dots*). Abbreviations—BST, bed nucleus of the stria terminalis; Ce, dorsal amygdala in the region of the central nucleus; FDR, false discovery rate; HDI, highest density interval; *t*, Student’s *t*; WB, whole-brain-corrected.

As shown in **Figure 5b**, the same general pattern was evident for skin conductance, an objective psychophysiological index of arousal (*Valence: F*(1,216)=790.55, *p*<0.001; *Certainty: F*(1,216)=138.95, *p*<0.001; *Valence × Certainty: F*(1,216)=661.63, *p*<0.001; *Threat, Uncertain vs. Certain: F*(1,216)=455.78, *p*<0.001; *Safety, Uncertain vs. Certain: F*(1,216)=270.03, *p*<0.001). These observations confirm the validity of the threat-anticipation paradigm as an experimental probe of fear and anxiety, consistent with prior work (<u>Hur et al., 2020b</u>; <u>Chavanne and Robinson, 2021</u>; <u>Hur et al., 2022</u>; <u>Kim et al., 2023</u>).

### N/NE amplifies distress evoked by the threat-anticipation paradigm

Exploratory analyses demonstrated that individuals with a more negative disposition experienced pervasively elevated distress across the four conditions of the threat-anticipation paradigm—both aversive and benign (*F*(1,218)=33.57, *p*<0.001)—and modestly potentiated reactivity to the anticipation of Threat compared to Safety, and to the anticipation of uncertain compared to certain outcomes (*N/NE × Valence: F*(1,218)=6.38, *p*=0.01; *N/NE × Certainty: F*(1,218)=6.03, *p*=0.02; **Figure 6**). No other moderator effects were significant for either subjective distress or objective arousal (*p*>0.57). In short, individuals with a more negative disposition show a mixture of indiscriminately elevated (‘overgeneralized’), threat-potentiated, and uncertainty-potentiated anticipatory distress, in broad accord with prior work (<u>Grupe and Nitschke, 2013</u>; <u>Shackman et al., 2016</u>).

**Figure 6.**
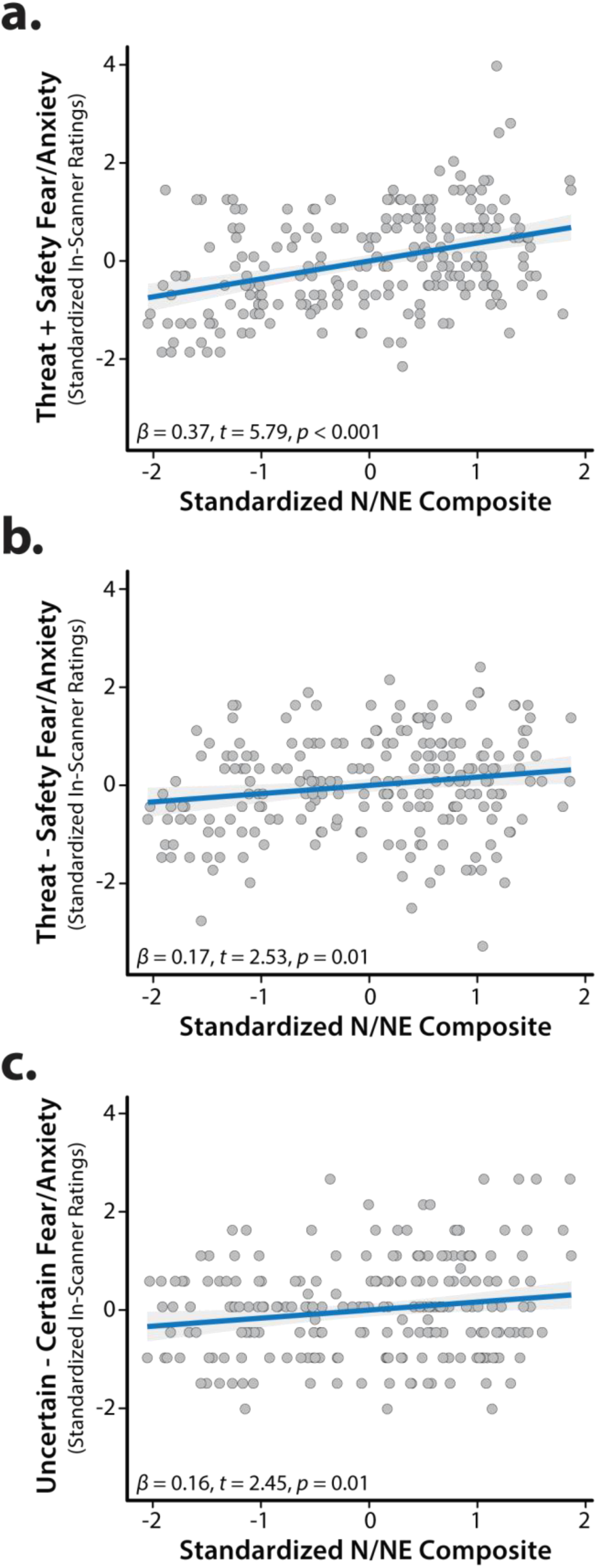
N/NE amplifies subjective fear and anxiety elicited by the threat-anticipation paradigm. Individuals with a more negative disposition exhibited a mixture of **(a)** indiscriminately elevated (‘overgeneralized’) distress across threat and safety trials, **(b)** potentiated distress when anticipating threat relative to safety, and **(c)** potentiated distress when anticipating temporally uncertain reinforcers. Scatterplots depict standard (ordinary least squares) regression estimates. Gray bands indicate 95% confidence intervals. Abbreviations—N/NE, neuroticism/negative emotionality.

### Threat anticipation and threat perception robustly recruit the EAc

We used whole-brain voxelwise GLMs to confirm that the threat-anticipation and threat-perception (emotional-faces) paradigms had the intended consequences on brain function. As expected, the anticipation of threat, uncertain threat, and certain threat significantly recruited both the BST and the dorsal amygdala in the region of the Ce (FDR *q*<.05, whole-brain corrected; **Figure 5c**). Beyond the EAc, each of these contrasts was associated with significant activation across a widely distributed network of regions previously implicated in the expression and regulation of human fear and anxiety (<u>Shackman and Fox, 2021</u>), including the midcingulate cortex, anterior insula/frontal operculum, dorsolateral prefrontal cortex, and periaqueductal grey (**Supplemental Tables 5-1, 5-2,** and **5-3;** https://osf.io/w5cdk). Analyses focused on the acute presentation of threat-related faces revealed significant activation in the region of the BST and the dorsal amygdala, consistent with prior work (**Figure 5c** and **Supplemental Table 5-4;** https://osf.io/w5cdk) (<u>Fox and Shackman, 2019</u>). Together, these observations demonstrate that our neuroimaging approach provides a robust probe of EAc function.

### BST and Ce are similarly sensitive to certain-versus-uncertain threat anticipation

As a precursor to hypothesis testing, we used a series of *t*-tests to confirm that the anatomically defined BST and Ce ROIs are recruited by the threat-anticipation and threat-perception (emotional-faces) tasks. Consistent with the voxelwise results (**Figure 5c**), both ROIs did, in fact, show nominally significant activation (*t*s(219)>3.10, *p*s<0.002, uncorrected; **Figure 7**). A standard 2 (*Region:* BST, Ce) × 2 (*Certainty:* Certain, Uncertain) GLM was used to explore potential regional differences in sensitivity to threat certainty. As shown in **Figure 7a** and detailed in **Supplemental Tables 7-1 and 7-2** (https://osf.io/w5cdk), although the BST was more strongly activated, on average, by anticipated threat (*Region: F*(1,219)=9.36, *p*=0.002), neither the main effect of Certainty nor the Region × Certainty (‘double-dissociation’) interaction was significant (*Certainty*: *F*(1,219)=0.19, *p*=0.66; *Region × Certainty*: *F*(1,219)=0.85, *p*=0.36). Similarly, the direct contrast of the two threat-anticipation conditions was not significant in either region (*BST*: *t*(219)=-0.82, *p*=0.41; *Ce*: *t*(219)=0.36, *p*=0.72). Collectively, these observations indicate the BST and Ce are similarly sensitive to temporally certain and uncertain threat, extending prior work by our group (<u>Hur et al., 2020b</u>).

**Figure 7.**
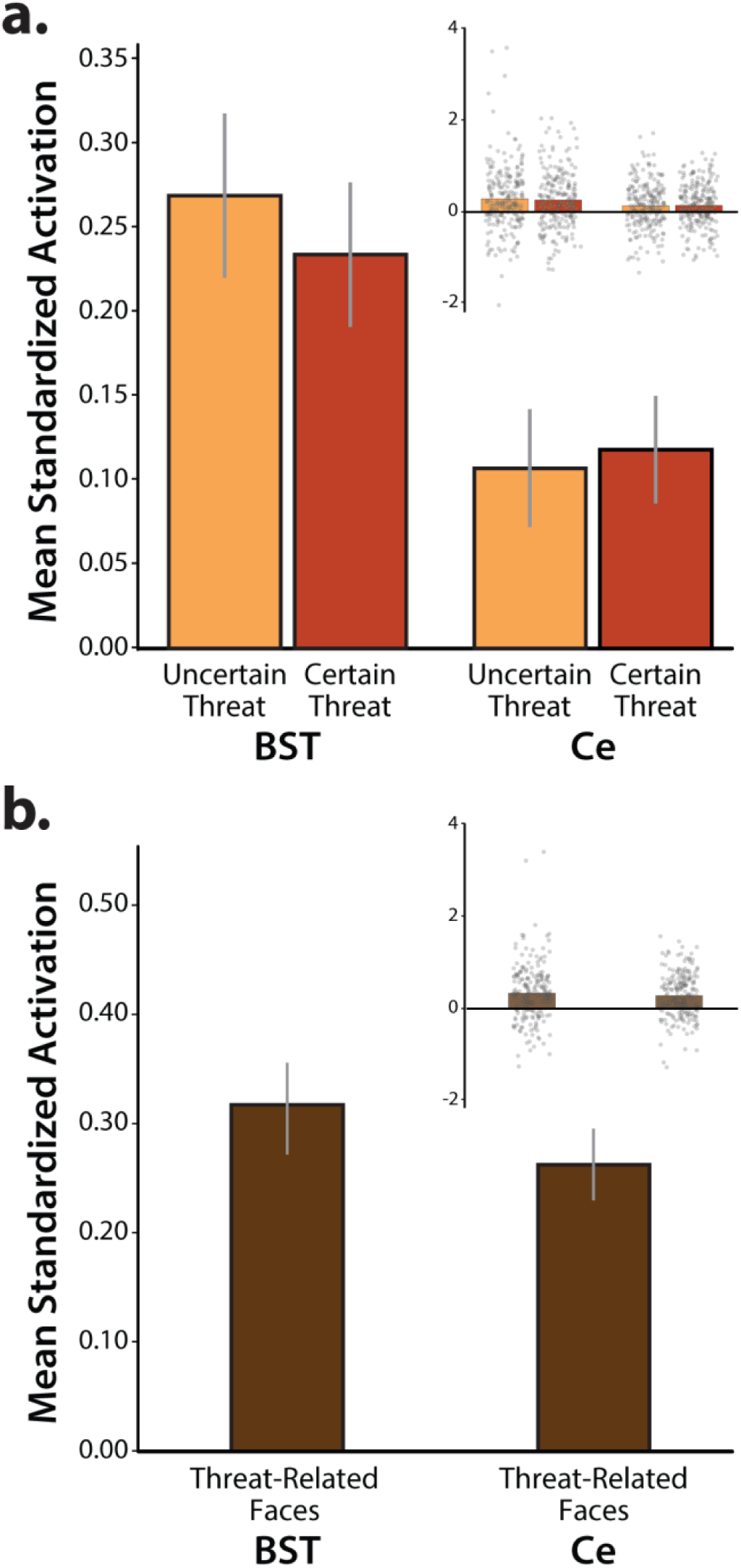
EAc ROI reactivity. The anatomically defined Ce and BST ROIs showed nominally significant activation during the **(a)** anticipation of threat and **(b)** presentation of threat-related faces (*p*s<0.003). Relative to the Ce, the BST was more strongly recruited during periods of threat anticipation (*p*=0.002). Neither the main effect of Certainty nor the Region × Certainty interaction was significant (*p*>0.18). Likewise, the direct contrast of certain-versus-uncertain threat was not significant in either ROI *(p*>0.40). For additional details, see **Supplemental Tables 7-1** and **7-2** (https://osf.io/w5cdk). Bars depict mean standardized ROI activation relative to the respective baseline conditions for each ROI (spatially unsmoothed data). Whiskers depict standard errors. Inset panels depict individual participants (*gray dots*). Abbreviations—BST, bed nucleus of the stria terminalis ROI; Ce, central nucleus of the amygdala ROI; ROI, region of interest.

### N/NE is associated with heightened BST activation during uncertain-threat anticipation

As shown schematically in **Figure 1**, cross-validated robust regressions were used to test the hypothesis that individuals with a more negative disposition will show exaggerated recruitment of the EAc (BST and/or Ce) during threat anticipation, and determine whether this association is more evident when the timing of threat encounters is uncertain. Results revealed that general EAc threat reactivity—aggregating across the anticipation of certain and uncertain aversive stimulation—was unrelated to individual differences in N/NE (*BST: β*=0.12, *t*(218)=1.57, *p*=0.12; *Ce: β*=0.02, *t*(218)=0.32, *p*=0.75; **Figures 8a-b**). Prior work suggests that relations between N/NE and EAc function may be magnified when threat encounters are uncertain in their timing or likelihood (<u>Somerville et al., 2010</u>). To test this, we computed robust regressions between N/NE and EAc reactivity to *uncertain* threat, separately for each EAc region. To clarify specificity, models controlled for activation during *certain* threat anticipation. Results demonstrated that heightened BST activation during uncertain-threat anticipation was significantly and selectively associated with trait-like variation in N/NE (*Uncertain: β*=0.24, *t*(217)=2.61, *p*=0.01; *Certain: β*=-0.09, *t*(217)=-1.00, *p*=0.32; **Figures 8c-d**). Follow-up analyses indicated that this association was significantly stronger for uncertain threat (*t_Hotelling_*(217)=2.42, *p*=0.02). Leveraging a simpler bivariate model, we estimated that BST reactivity to uncertain-threat anticipation explained, on average, 5.1% of the variance in N/NE in out-of-sample test folds (*β*=0.19, *t*(218)=2.59, *p*=0.01, cross-validated *R^2^*=0.051). BST reactivity was unrelated to individual differences in task-related distress (|*t*|<1.00, *p*>0.31).

**Figure 8.**
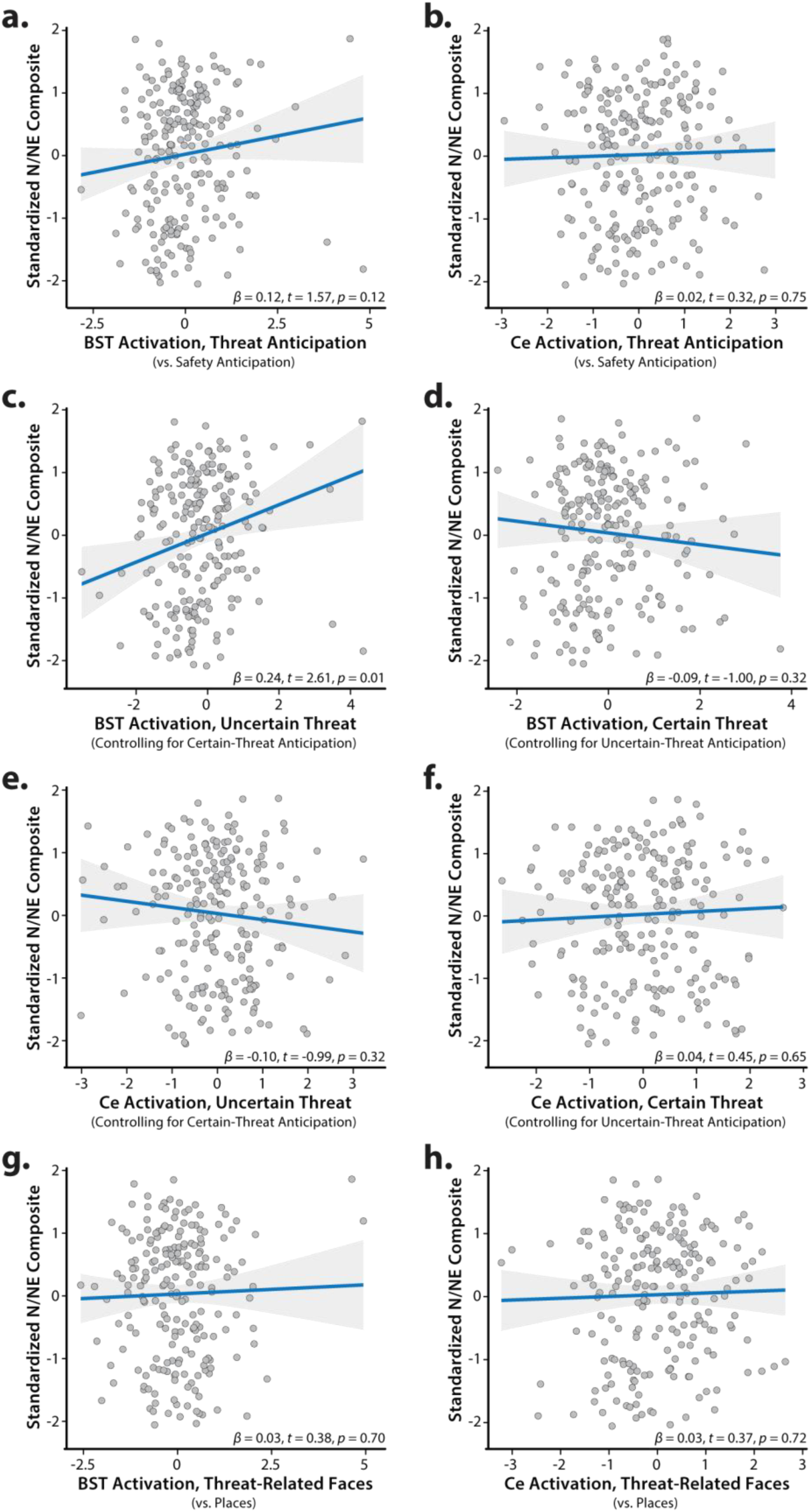
Individual differences in N/NE are associated with heightened BST activation during the uncertain anticipation of a genuinely distressing threat. Scatterplots depict the robust association (*blue line*) between N/NE (standardized multi-scale/multi-occasion composite) and each task-related neuroimaging metric for the BST and Ce ROIs. Panels *c*-*f* depict partial correlations. Activation estimates for the anticipation of certain/uncertain threat and the presentation of threat-related faces reflect differences relative to the implicit baseline. Gray bands indicate 95% confidence intervals. Abbreviations— BST, bed nucleus of the stria terminalis ROI; Ce, central nucleus of the amygdala ROI; N/NE, neuroticism/negative emotionality.

In contrast to the BST, Ce reactivity to the threat-anticipation task was unrelated to variation in N/NE, regardless of threat certainty (*Uncertain: β*=-0.10, *t*(217)=-0.99, *p*=0.32; *Certain: β*=0.04, *t*(217)=0.45, *p*=0.65; **Figures 8e-f**). EAc reactivity to the threat-perception (emotional-faces) task was also unrelated to N/NE (*BST: β*=0.03, *t*(211)=0.38, *p*=0.70; *Ce: β*=0.03, *t*(211)=0.37, *p*=0.72; **Figures 8g-h**). Consistent with these nil effects, the association between BST reactivity to uncertain threat and N/NE remained significant in models that controlled for either Ce reactivity to uncertain-threat anticipation or BST reactivity to threat-related faces (*t*>2.59, *p*<0.02). The association between N/NE and BST reactivity to uncertain threat was significantly greater than that obtained for Ce reactivity to uncertain threat (*t_Hotelling_*(217)=2.94, *p*=0.004), but did not differ from that obtained for BST reactivity to threat-related faces (*t_Hotelling_*(206)=1.57, *p*=0.12). In sum, across the regions and tasks considered here, individual differences in N/NE are uniquely associated with heightened BST activation during the uncertain anticipation of a genuinely distressing threat (**Figure 9**).

**Figure 9.**
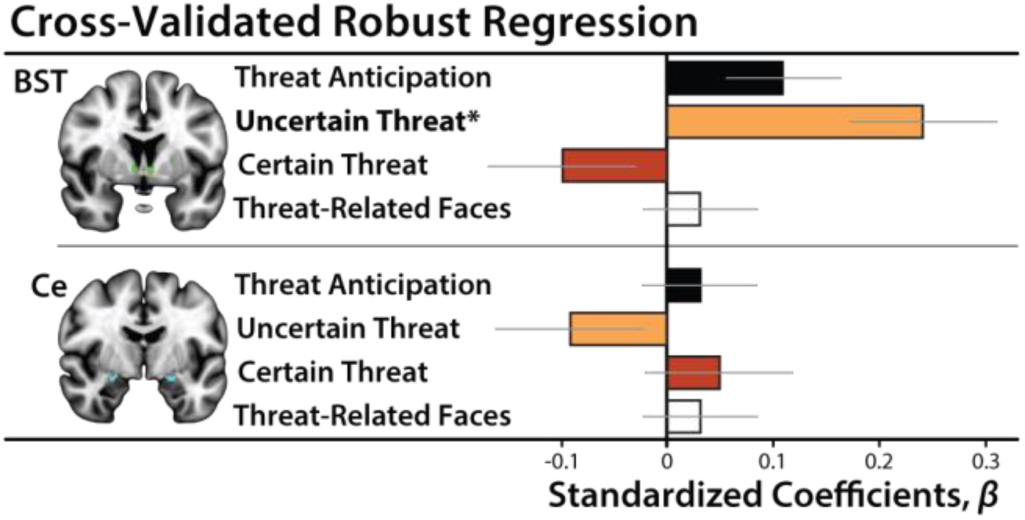
Summary of robust regression models. Heightened BST reactivity to uncertain-threat anticipation (*orange*) was associated with elevated levels of N/NE (*p*=0.01). Other associations were not significant (*p*>0.11). Bars depict standardized coefficients for each robust regression model. Whiskers indicate standard errors. Significant associations are marked by an asterisk. Abbreviations—BST, bed nucleus of the stria terminalis ROI; Ce, central nucleus of the amygdala ROI.

### BST reactivity to uncertain threat is broadly associated with the ‘internalizing’ facets of N/NE

Epidemiological, psychiatric, and biological studies typically focus on coarse ‘broadband’ measures of N/NE (<u>Shackman et al., 2016</u>; <u>Hur et al., 2019</u>). Yet it is clear that N/NE is a complex phenotype that subsumes several narrower traits—including dispositional anxiety, depression/sadness, and emotional volatility (<u>Caspi et al., 2005</u>; <u>Soto and John, 2017</u>; <u>Kalokerinos et al., 2020</u>)—each characterized by a mixture of shared and unique psychological associations and biological correlates (<u>Thorp et al., 2021</u>; <u>Klein-Flügge et al., 2022</u>; <u>Watson et al., 2022</u>; <u>Khoo et al., 2023</u>). While our composite N/NE instrument has many psychometric strengths (**Figure 1**), it cannot address which of these facets is most relevant to BST function (<u>McCrae, 2015</u>). To do so, we leveraged the revised Big Five Inventory (BFI-2), a well-established, hierarchically organized scale that was expressly constructed to enable rigorous facet-level analyses (<u>Soto and John, 2017</u>). The BFI-2 was administered at the baseline (T2) and 6-month follow-up (T3) sessions (**Figure 1**). Paralleling the approach used for broadband N/NE, facet scores were averaged across assessments to minimize occasion-specific fluctuations (‘noise’). Cross-validated robust GLMs were used to quantify associations between BST reactivity to uncertain-threat anticipation and each facet of N/NE, while controlling for BST reactivity to certain threat. Results revealed significant associations with dispositional Anxiety and Depression/Sadness, but not Emotional Volatility (*Anxiety: β*=0.20, *t*(217)=2.19, *p*=0.03; *Depression/Sadness: β*=0.22, *t*(217)=2.45, *p*=0.02; *Volatility: β*=0.10, *t*(217)=1.10, *p*=0.27). Consistent with the broadband results, BST reactivity to certain-threat anticipation was unrelated to the three narrow traits (*p*>0.21). In cross-validated bivariate models, variation in BST reactivity to uncertain-threat anticipation explained an average of ∼4% of the variance in the Anxiety and Depression/Sadness facets of N/NE in out-of-sample test data (*Anxiety: β*=0.15, *t*(218)=2.13, *p*=0.04, cross-validated *R^2^*=0.041; *Depression/Sadness: β*=0.16, *t*(218)=2.17, *p*=0.03, cross-validated *R^2^*=0.040). While the BST is often conceptualized as playing a central role in anxiety-related states, traits, and disorders (<u>Fox and Shackman, 2019</u>; <u>Shackman and Fox, 2021</u>; <u>Grogans et al., 2023</u>), these findings demonstrate that heightened BST reactivity to uncertain threat is more broadly associated with the ‘internalizing’ facets of N/NE (Anxiety and Depression/Sadness). They also indicate that our major conclusions generalize across questionnaire instruments.

### EAc reactivity to the threat-anticipation and threat-perception tasks shows negligible convergence

It is often assumed that different experimental manipulations of ‘threat’ are fungible probes of individual differences in circuit function (i.e., threat-of-shock ≈ threat-related faces). Yet this tacit assumption of convergent validity has never been tested in a larger sample or in the BST (<u>Villalta-Gil et al., 2017</u>). As shown in **Table 2**, robust GLMs revealed negligible associations between BST reactivity to the threat-anticipation and threat-perception (emotional-faces) tasks (*p*>0.06). The same pattern of null associations was evident for the Ce (*p*>0.11). The absence of robust cross-task correlations raises important questions about the equivalence of two popular fMRI threat tasks—one centered on the cued anticipation of aversive stimulation, the other focused on the perception of angry and fearful facial expressions—and caution against relying exclusively on emotional-face paradigms to probe individual differences in threat-related EAc function (<u>Grogans et al., 2022</u>).

**Table 2.** Convergent validity of EAc reactivity to the threat-anticipation and threat-perception paradigms (anatomically defined ROIs; spatially unsmoothed data).

| ROI | Contrast A<sup>a</sup> | Contrast B<sup>a</sup> | Robust $\beta$ | $t(207)^b$ | Nominal $p$ |
| --- | --- | --- | --- | --- | --- |
| BST | Threat-Related Faces | Threat Anticipation | 0.11 | 1.80 | 0.07 |
|  |  | Uncertain-Threat Anticipation | 0.02 | 0.42 | 0.68 |
|  |  | Certain-Threat Anticipation | 0.12 | 1.68 | 0.10 |
| Ce | Threat-Related Faces | Threat Anticipation | -0.01 | -0.20 | 0.84 |
|  |  | Uncertain-Threat Anticipation | -0.10 | -1.47 | 0.14 |
|  |  | Certain-Threat Anticipation | -0.11 | -1.54 | 0.12 |

<sup>a</sup> Relative to their respective reference conditions. <sup>b</sup> A total of 209 participants provided usable data for both paradigms. Abbreviations—BST, bed nucleus of the stria terminalis; Ce, dorsal amygdala in the region of the central nucleus; EAc, central extended amygdala; ROI, region of interest.

## DISCUSSION (1,574/1,500)

Our results show that the threat-anticipation paradigm elicited robust distress and arousal, confirming its validity as a N/NE-relevant challenge (**Figure 5**). Fearful and anxious feelings were more intense and indiscriminate among high-N/NE individuals, with elevated distress apparent even while waiting for benign stimulation (**Figure 6**). Both threat paradigms strongly recruited the EAc, indicating that our approach is appropriate for probing its function (**Figures 5** and **7**). Cross-validated robust regressions showed that N/NE is associated with heightened BST activation during the temporally uncertain anticipation of threat (**Figures 8** and **9**). In contrast, N/NE was unrelated to BST activation during certain-threat anticipation, Ce activation during either type of threat anticipation, or BST/Ce reactivity to threat-related faces. While the BST is often conceptualized in terms of anxiety, our results suggest that it is broadly associated with the internalizing facets of N/NE, including the tendency to experience sadness and depression. While it is often assumed that different threat paradigms are interchangeable measures of individual differences in EAc function, our results revealed negligible cross-task associations in the EAc (**Table 2**).

Our results show that N/NE is associated with heightened BST reactivity to uncertain-threat anticipation, extending prior work (<u>Shackman et al., 2016</u>). Differences in N/NE are heritable and work in monkeys suggests that variation in BST reactivity to uncertain threat is particularly relevant to the heritable component of N/NE (<u>Fox et al., 2015b</u>), although we acknowledge that it remains unknown whether this association is specific to uncertain-threat anticipation or reflects a broader hyper-sensitivity to uncertainty (<u>Grupe and Nitschke, 2013</u>). Anatomical tracing studies indicate that the BST is poised to assemble behavioral, psychophysiological, and neuroendocrine signs of negative affect via projections to downstream effectors (<u>Fox et al., 2015a</u>). Collectively, these observations reinforce the hypothesis that the BST is an evolutionarily conserved component of the distributed neural system governing trait-like individual differences in threat reactivity. A key challenge will be to clarify causation. Although the mechanistic contribution of the BST to dispositional fear and anxiety has yet to be explored in primates, work in rodents shows that the BST exerts bi-directional control over defensive behaviors triggered by uncertain-threat anticipation (<u>Jennings et al., 2013</u>; <u>Kim et al., 2013</u>; <u>Glangetas et al., 2017</u>; <u>Ahrens et al., 2018</u>; <u>Mazzone et al., 2018</u>). Among humans, N/NE is stable, but can change in response to experience, providing opportunities for understanding causation (<u>Stieger et al., 2021</u>; <u>Zemestani et al., 2022</u>). In a comprehensive meta-analysis, Roberts and colleagues documented substantial N/NE reductions following treatment for internalizing disorders (<u>Roberts et al., 2017</u>). It will be fruitful to determine whether this reflects dampened BST reactivity to uncertain threat, and whether it depends on treatment modality (i.e., psychosocial vs. pharmacological).

Elevated N/NE is a prominent risk factor for pathological anxiety and depression (<u>Shackman et al., 2016</u>; <u>Toenders et al., 2022</u>). Co-morbidity between anxiety and depression disorders is rampant and ∼75% of patients with major depression exhibit significant anxiety symptoms (<u>Hasin et al., 2018</u>; <u>Caspi et al., 2020</u>). Furthermore, anxiety, depression, and N/NE show robust genetic correlations and parallel pharmacological effects (<u>Barlow et al., 2013</u>; <u>Levey et al., 2020</u>; <u>Thorp et al., 2021</u>). Although these findings suggest a common neurobiological substrate (<u>Barlow et al., 2013</u>), identifying the relevant circuitry has proven challenging (<u>Grogans et al., 2022</u>). Our observation that heightened BST reactivity to uncertain threat is associated with both the anxious and the depressive facets of N/NE suggests that alterations in BST threat-reactivity might contribute to this still-enigmatic shared substrate. While this hypothesis remains untested, the available evidence is supportive. A recent meta-analysis demonstrated that the BST is hyper-reactive to unpleasant emotional challenges among individuals with anxiety disorders (<u>Chavanne and Robinson, 2021</u>). Both anxiety and depression are often treated via chronic administration of selective serotonin reuptake inhibitors (SSRIs) or, in the case of anxiety, acute administration of benzodiazepines. Both treatments have been shown to disproportionately reduce defensive responses to uncertain threat (relative to certain threat) in humans and rats, and to dampen BST reactivity to uncertain threat in rats (<u>Bechtholt et al., 2008</u>; <u>Grillon and Ernst, 2020</u>). A key avenue for future research will be to use prospective-longitudinal data to test whether exaggerated BST activation during uncertain-threat anticipation increases the likelihood of future anxiety and depression.

Our results have implications for the design and interpretation of neuroimaging studies of psychiatric risk, disease, and treatment. Most of this work relies on emotional-face tasks as the lone probe of fear, anxiety, and related ‘Negative Valence Systems.’ Yet our results indicate that Ce and BST reactivity to threat-related faces is unrelated to the risk-conferring N/NE phenotype. These null effects are not unprecedented. Three other well-powered studies failed to detect associations between amygdala face reactivity and N/NE (<u>MacDuffie et al., 2019</u>; <u>Silverman et al., 2019</u>; <u>West et al., 2021</u>). Our results also make it clear that the acute perception of threat-related faces and the anticipation of aversive stimulation are statistically distinct assays of individual differences in EAc function, consistent with prior work in a much smaller sample (<u>Villalta-Gil et al., 2017</u>). These observations highlight the hazards of continuing to rely on a limited number of ‘workhorse’ neuroimaging paradigms to understand and predict emotion, temperament, and psychopathology (<u>Grogans et al., 2023</u>). They also caution against muddling the distinction between the perception of threat-related cues and the actual experience of fear and anxiety, a practice that has become routine in therapeutics research (<u>Kwako et al., 2015</u>; <u>Schwandt et al., 2016</u>; <u>Paulus et al., 2021</u>; <u>Bloomfield et al., 2022</u>).

Our results indicate that BST reactivity to uncertain threat predicts 5.1% of the variance in N/NE in out-of-sample data. The magnitude of this association—while too modest for practical applications—appears plausible, given the complexity of the N/NE phenotype and BST function—and compares favorably with other psychiatrically relevant associations, including prospective associations between striatal reward reactivity and depression (1%) and amygdala face reactivity and internalizing symptoms (2.7%) (<u>Grogans et al., 2022</u>). It exceeds the predictive performance (1.5-4.2%) of polygenic scores derived from large-scale GWAS of N/NE (<u>Nagel et al., 2018</u>; <u>Baselmans et al., 2019</u>). Notably, the magnitude of our BST ‘hit’ is half that of the cross-validated association reported for dorsal-attention network activation during a working-memory paradigm and general-cognitive ability (11.6%), suggesting that it is not unrealistically large (<u>Marek et al., 2022</u>). From a mechanistic perspective, the small-but-reliable hits uncovered by adequately powered association studies are useful for prioritizing targets for perturbation studies. This reflects the fact that modest brain-behavior associations do not preclude substantially larger effects with targeted interventions (<u>Shackman and Fox, 2018</u>). Indeed, EAc damage can have dramatic consequences for fear-and anxiety-related states and traits (<u>Fox and Shackman, 2019</u>).

Our results also have implications for psychological theories of N/NE. Most theories are rooted in the idea that N/NE reflects hyper-sensitivity to threat, amplifying emotional responses when stressors are encountered (<u>Spielberger, 1966</u>; <u>Eysenck, 1967</u>; <u>Kagan et al., 1988</u>). Indeed, we found that high-N/NE individuals experienced potentiated distress when anticipating threat relative to safety (*R^2^*=2.9%; **Figure 6b**). Yet high-N/NE individuals also reported heightened distress when anticipating outcomes—whether aversive or benign—that were simply uncertain (*R^2^*=2.6%; **Figure 6c**), consistent with models emphasizing the centrality of uncertainty to anxiety (<u>Grupe and Nitschke, 2013</u>). But by far the strongest effect of N/NE was indiscriminately elevated distress across threat and safety trials (*R^2^*=13.7%; **Figure 6a**). This finding is consistent with other work using threat-anticipation paradigms (<u>Sep et al., 2019</u>; <u>Stegmann et al., 2019</u>) and extends research focused on more naturalistic provocations (e.g., aversive films) and smartphone measures of real-world experience (<u>Shackman et al., 2016</u>; <u>Hur et al., 2022</u>). Collectively, these observations reinforce models that emphasize the importance of pervasive, contextually inappropriate negative affect (<u>Shackman et al., 2016</u>). From a psychiatric perspective, overgeneralized responses to threat are particularly noteworthy because they promote avoidance, a key feature of pathological anxiety; distinguish patients from controls; and confer risk for future psychopathology (<u>Shackman et al., 2016</u>; <u>Hunt et al., 2017</u>).

Clearly, important challenges remain. First, our study was focused on an ethnoracially diverse sample of emerging adults. It will be useful to expand this to encompass more representative samples. Second, although our results highlight the importance of the BST, N/NE is a complex, multidimensional phenotype that undoubtedly reflects multiple regions and networks. It will be important to understand how interactions between the BST and other regions implicated in the control of negative affect support variation in N/NE. Third, the absence of reward trials precludes strong claims about valence. While unlikely, similar associations might be evident for uncertain-reward anticipation. Fourth, BST function was unrelated to task-related distress, raising questions about the precise mechanisms linking it to variation in N/NE. This null association might reflect an artifact of our sparse distress-sampling protocol (16.7% trials) or it could be that BST reactivity to uncertain-threat anticipation only indirectly influences conscious feelings (<u>Taschereau-Dumouchel et al., 2022</u>). Moving forward, it will be useful to collect trial-by-trial fear/anxiety ratings, enabling formal mediation analyses, and to assess relations with real-world distress.

Elevated N/NE is associated with a multitude of practically important outcomes, yet the underlying neurobiology has remained unclear. Our observations show that N/NE is associated with heightened activation during the anticipation of an uncertain, genuinely distressing threat in the BST, but not in the Ce. EAc reactivity to threat-related faces was also unrelated to N/NE. These observations provide a neurobiological framework for conceptualizing N/NE and set the stage for more ambitious prospective-longitudinal and mechanistic studies. A comparatively large, diverse, and carefully phenotyped sample and a pre-registered, best-practices approach enhance confidence in these results.

## ACKNOWLEDGMENTS

Authors acknowledge assistance and critical feedback from three anonymous reviewers, A. Antonacci, L. Friedman, J. Furcolo, M. Gamer, C. Grubb, R. Hum, C. Kaplan, T. Kashdan, J. Kuang, C. Lejuez, D. Limon, B. Nacewicz, L. Pessoa, S. Rose, J. Swayambunathan, A. Vogel, J. Wedlock, members of the Affective and Translational Neuroscience laboratory, the staff of the Maryland Neuroimaging Center, and the Office of the Registrar at the University of Maryland. This work was partially supported by the California National Primate Center; National Institutes of Health (AA030042, DA040717, MH018921, MH107444, MH121409, MH121735, MH128336, MH129851, OD011107, MH131264, MH132280); National Research Foundation of Korea (2021R1F1A1063385 and 2021S1A5A2A03070229); University of California, Davis; University of Maryland; and Yonsei Signature Research Cluster Program (2021-22-0005). Authors declare no conflicts of interest.

## AUTHOR CONTRIBUTIONS

A.J.S., K.A.D., and J.F.S. designed the overall study. S.E.G. and A.J.S. envisioned the present project. J.F.S. and

A.J.S. developed and optimized the imaging paradigm. K.A.D. managed data collection and study administration. K.A.D., J.F.S., A.S.A, S.I., and R.M.T. collected data. J.F.S. and M.K. developed data processing and analytic software for imaging analyses. J.H., J.F.S., H.C.K., and R.M.T. processed imaging data. S.E.G., J.F.S., and A.J.S. analyzed data. S.E.G., M.G.B., and A.J.S. developed the analytic strategy. S.E.G., A.S.F., J.F.S., and A.J.S. interpreted data. S.E.G. and A.J.S. wrote the paper. S.E.G. created figures and tables. A.J.S. funded and supervised all aspects of the study. All authors contributed to reviewing and revising the paper and approved the final version.

## PREREGISTRATION

Our general approach and hypotheses were preregistered (https://osf.io/wzhdm).

## RESOURCE SHARING

Raw data and select materials are publicly available at the National Institute of Mental Health Data Archive (https://nda.nih.gov/edit_collection.html?id=2447). Processed data and supplementary tables (e.g., cluster tables) are available at the Open Science Framework (https://osf.io/w5cdk). Key neuroimaging maps are available at NeuroVault (https://neurovault.org/collections/13109).

