## Extended Data Table 5-1 for "Neuroticism/negative emotionality is associated with increased reactivity to uncertain threat in the bed nucleus of the stria terminalis, not the amygdala" (from another version of this preprint: version 5, 2024-04-21)

**Extended Data Table 5-1.** Descriptive statistics for clusters and local extrema showing greater activation during the anticipation of Threat relative to Safety (FDR *q*<0.05, whole-brain corrected; spatially unsmoothed data).

|  | | **mm^3^** | ***t*** | **x** | **y** | **z** |
| --- | --- | --- | --- | --- | --- | --- |
|  | **Cluster 1** | 666,016 |  |  |  |  |
|  | L Frontal Operculum Cortex |  | 13.64 | -32 | 20 | 10 |
|  | L Frontal Orbital Cortex |  | 11.77 | -30 | 28 | 0 |
|  | L Paracingulate Gyrus |  | 11.56 | -8 | 12 | 38 |
|  | L Cingulate Gyrus, anterior division |  | 11.50 | -6 | 10 | 40 |
|  | L Central Opercular Cortex |  | 11.10 | -42 | 8 | 2 |
|  | L Caudate |  | 11.05 | -8 | 0 | 8 |
|  | L Putamen |  | 10.88 | -20 | 10 | -2 |
|  | L Temporal Occipital Fusiform Cortex |  | 10.67 | -26 | -60 | -16 |
|  | L Brain Stem |  | 10.30 | -2 | -28 | -2 |
|  | L Superior Frontal Gyrus |  | 10.25 | -14 | -2 | 66 |
|  | L Precentral Gyrus |  | 10.07 | -32 | -6 | 50 |
|  | L Juxtapositional Lobule Cortex |  | 10.05 | -2 | 6 | 46 |
|  | L Cingulate Gyrus, posterior division |  | 10.04 | -14 | -28 | 38 |
|  | L Thalamus |  | 10.01 | -8 | -4 | 12 |
|  | L Occipital Fusiform Gyrus |  | 9.48 | -30 | -68 | -18 |
|  | L Supramarginal Gyrus, posterior division |  | 9.45 | -56 | -50 | 38 |
|  | L Bed Nucleus of the Stria Terminalis |  | 9.43 | -8 | 2 | 4 |
|  | L Middle Frontal Gyrus |  | 9.16 | -42 | -2 | 60 |
|  | L Supramarginal Gyrus, anterior division |  | 8.89 | -54 | -38 | 32 |
|  | L Inferior Frontal Gyrus, pars opercularis |  | 8.85 | -54 | 10 | 10 |
|  | L Lingual Gyrus |  | 8.85 | -4 | -74 | -12 |
|  | L Frontal Pole |  | 8.83 | -36 | 44 | 30 |
|  | L Bed Nucleus of the Stria Terminalis |  | 8.55 | -6 | 4 | 0 |
|  | L Parietal Operculum Cortex |  | 7.71 | -56 | -34 | 24 |
|  | L Angular Gyrus |  | 7.67 | -56 | -56 | 46 |
|  | L Superior Parietal Lobule |  | 7.55 | -18 | -52 | 64 |
|  | L Precuneus Cortex |  | 7.49 | -10 | -78 | 40 |
|  | L Lateral Occipital Cortex, superior division |  | 7.35 | -34 | -60 | 60 |
|  | L Postcentral Gyrus |  | 7.27 | -42 | -32 | 44 |
|  | L Cuneal Cortex |  | 6.34 | -16 | -78 | 34 |
|  | L Occipital Pole |  | 5.90 | -20 | -96 | 10 |
|  | L Hippocampus |  | 5.56 | -34 | -28 | -8 |
|  | L Insular Cortex |  | 5.55 | -34 | 18 | -4 |
|  | L Lateral Occipital Cortex, inferior division |  | 5.50 | -56 | -64 | 10 |
|  | L Pallidum |  | 5.50 | -20 | -10 | -4 |
|  | L Inferior Frontal Gyrus, pars triangularis |  | 5.46 | -54 | 22 | 6 |
|  | L Inferior Temporal Gyrus, temporooccipital part |  | 5.46 | -42 | -56 | -10 |
|  | L Heschls Gyrus (includes H1 and H2) |  | 5.31 | -38 | -28 | 12 |
|  | L Middle Temporal Gyrus, temporooccipital part |  | 5.13 | -62 | -56 | 6 |
|  | L Planum Polare |  | 5.06 | -40 | -14 | -8 |
|  | L Temporal Fusiform Cortex, posterior division |  | 5.01 | -30 | -32 | -28 |
|  | L Amygdala |  | 4.74 | -24 | -14 | -14 |
|  | L Superior Temporal Gyrus, posterior division |  | 4.47 | -54 | -24 | -4 |
|  | L Middle Temporal Gyrus, posterior division |  | 4.44 | -48 | -28 | -6 |
|  | L Planum Temporale |  | 4.37 | -62 | -20 | 4 |
|  | L Intracalcarine Cortex |  | 4.34 | -18 | -66 | 6 |
|  | L Amygdala (*central nucleus*) |  | 4.14 | -24 | -12 | -12 |
|  | L Subcallosal Cortex |  | 4.12 | -12 | 16 | -14 |
|  | L Temporal Pole |  | 4.04 | -52 | 14 | -12 |
|  | L Accumbens |  | 3.65 | -6 | 12 | -6 |
|  | L Parahippocampal Gyrus, posterior division |  | 3.33 | -14 | -36 | -6 |
|  | L Superior Temporal Gyrus, anterior division |  | 3.12 | -54 | 2 | -14 |
|  | L Amygdala (*central nucleus*) |  | 2.92 | -22 | -6 | -12 |
|  | L Inferior Temporal Gyrus, posterior division |  | 2.34 | -50 | -38 | -16 |
|  | L Supracalcarine Cortex |  | 2.32 | -22 | -62 | 20 |
|  | R Cingulate Gyrus, anterior division |  | 14.08 | 10 | 12 | 38 |
|  | R Paracingulate Gyrus |  | 14.06 | 10 | 20 | 34 |
|  | R Frontal Operculum Cortex |  | 13.18 | 34 | 24 | 8 |
|  | R Bed Nucleus of the Stria Terminalis/Caudate |  | 12.87 | 10 | 2 | 8 |
|  | R Thalamus |  | 12.44 | 10 | -2 | 12 |
|  | R Precentral Gyrus |  | 12.08 | 42 | -2 | 46 |
|  | R Juxtapositional Lobule Cortex |  | 11.49 | 10 | 6 | 44 |
|  | R Putamen |  | 11.44 | 22 | 6 | 4 |
|  | R Brain Stem |  | 11.39 | 4 | -28 | -2 |
|  | R Supramarginal Gyrus, posterior division |  | 11.32 | 60 | -42 | 24 |
|  | R Central Opercular Cortex |  | 11.15 | 44 | 8 | 2 |
|  | R Superior Temporal Gyrus, posterior division |  | 10.61 | 48 | -24 | -4 |
|  | R Superior Frontal Gyrus |  | 10.41 | 14 | 10 | 62 |
|  | R Pallidum |  | 9.87 | 20 | 4 | 0 |
|  | R Superior Parietal Lobule |  | 9.69 | 24 | -46 | 62 |
|  | R Frontal Orbital Cortex |  | 9.57 | 34 | 22 | -8 |
|  | R Frontal Pole |  | 9.56 | 34 | 46 | 30 |
|  | R Bed Nucleus of the Stria Terminalis/Caudate |  | 9.51 | 8 | 6 | 0 |
|  | R Cingulate Gyrus, posterior division |  | 9.49 | 10 | -26 | 42 |
|  | R Parietal Operculum Cortex |  | 9.40 | 56 | -30 | 26 |
|  | R Supramarginal Gyrus, anterior division |  | 9.33 | 56 | -32 | 34 |
|  | R Angular Gyrus |  | 9.16 | 64 | -46 | 28 |
|  | R Precuneus Cortex |  | 8.97 | 12 | -54 | 54 |
|  | R Middle Frontal Gyrus |  | 8.84 | 52 | 8 | 42 |
|  | R Postcentral Gyrus |  | 8.43 | 48 | -28 | 50 |
|  | R Inferior Frontal Gyrus, pars triangularis |  | 8.05 | 50 | 22 | 2 |
|  | R Temporal Occipital Fusiform Cortex |  | 8.03 | 26 | -56 | -16 |
|  | R Middle Temporal Gyrus, posterior division |  | 7.83 | 52 | -30 | -4 |
|  | R Inferior Frontal Gyrus, pars opercularis |  | 7.67 | 54 | 12 | 4 |
|  | R Lateral Occipital Cortex, superior division |  | 7.50 | 18 | -74 | 40 |
|  | R Middle Temporal Gyrus, temporooccipital part |  | 6.94 | 56 | -40 | 4 |
|  | R Inferior Temporal Gyrus, temporooccipital part |  | 6.69 | 52 | -60 | -12 |
|  | R Occipital Fusiform Gyrus |  | 6.45 | 20 | -74 | -16 |
|  | R Occipital Pole |  | 6.41 | 28 | -94 | 12 |
|  | R Hippocampus |  | 6.38 | 34 | -28 | -6 |
|  | R Lateral Occipital Cortex, inferior division |  | 6.29 | 42 | -86 | -8 |
|  | R Lingual Gyrus |  | 6.27 | 0 | -74 | -8 |
|  | R Cuneal Cortex |  | 6.00 | 14 | -76 | 36 |
|  | R Insular Cortex |  | 5.32 | 42 | -2 | -12 |
|  | R Temporal Pole |  | 5.17 | 54 | 16 | -10 |
|  | R Accumbens |  | 4.77 | 6 | 10 | -6 |
|  | R Planum Polare |  | 4.50 | 44 | -8 | -6 |
|  | R Intracalcarine Cortex |  | 4.35 | 24 | -60 | 6 |
|  | R Amygdala (*central nucleus*) |  | 4.16 | 22 | -6 | -12 |
|  | R Temporal Fusiform Cortex, posterior division |  | 4.07 | 40 | -32 | -16 |
|  | R Inferior Temporal Gyrus, anterior division |  | 3.99 | 42 | -2 | -38 |
|  | R Supracalcarine Cortex |  | 3.89 | 22 | -64 | 14 |
|  | R Parahippocampal Gyrus, anterior division |  | 3.66 | 26 | -4 | -32 |
|  | R Planum Temporale |  | 3.54 | 40 | -28 | 12 |
|  | R Parahippocampal Gyrus, posterior division |  | 3.32 | 16 | -34 | -6 |
|  | R Heschls Gyrus (includes H1 and H2) |  | 2.95 | 48 | -8 | 4 |
|  | R Superior Temporal Gyrus, anterior division |  | 2.66 | 56 | 4 | -14 |
|  | R Temporal Fusiform Cortex, anterior division |  | 2.33 | 38 | -8 | -38 |
|  | **Cluster 2** | 1,368 |  |  |  |  |
|  | L Middle Temporal Gyrus, anterior division |  | 4.11 | -48 | -2 | -32 |
|  | L Inferior Temporal Gyrus, anterior division |  | 3.97 | -46 | 0 | -34 |
|  | L Inferior Temporal Gyrus, posterior division |  | 3.94 | -56 | -20 | -32 |
|  | L Temporal Pole |  | 3.61 | -42 | 10 | -34 |
|  | L Temporal Fusiform Cortex, anterior division |  | 3.44 | -34 | -6 | -34 |
|  | **Cluster 3** | 920 |  |  |  |  |
|  | R Frontal Pole |  | 4.64 | 26 | 52 | -12 |
|  | **Cluster 4** | 392 |  |  |  |  |
|  | R Frontal Pole |  | 3.88 | 30 | 68 | 2 |
|  | **Cluster 5** | 104 |  |  |  |  |
|  | L Frontal Pole |  | 3.22 | -24 | 66 | -6 |
|  | **Cluster 6** | 96 |  |  |  |  |
|  | R Occipital Pole |  | 3.88 | 6 | -96 | -8 |
|  | **Cluster 7** | 88 |  |  |  |  |
|  | R Paracingulate Gyrus |  | 3.14 | 12 | 46 | -2 |
|  | **Cluster 8** | 88 |  |  |  |  |
|  | L Insular Cortex |  | 4.28 | -42 | -14 | 2 |
|  | **Cluster 9** | 80 |  |  |  |  |
|  | L Hippocampus |  | 3.51 | -30 | -14 | -24 |
|  | L Parahippocampal Gyrus, anterior division |  | 2.44 | -30 | -10 | -30 |
|  | **Cluster 10** | 80 |  |  |  |  |
|  | R Frontal Pole |  | 2.40 | 18 | 44 | 44 |
|  | **Cluster 11** | 64 |  |  |  |  |
|  | L Temporal Pole |  | 3.21 | -44 | 16 | -26 |
|  | **Cluster 12** | 64 |  |  |  |  |
|  | L Temporal Pole |  | 4.57 | -50 | 16 | -18 |
|  | **Cluster 13** | 56 |  |  |  |  |
|  | L Middle Temporal Gyrus, posterior division |  | 2.61 | -66 | -34 | -18 |
|  | **Cluster 14** | 56 |  |  |  |  |
|  | L Frontal Pole |  | 2.95 | -46 | 38 | 4 |
|  | L Inferior Frontal Gyrus, pars triangularis |  | 2.32 | -52 | 36 | 6 |
|  | **Cluster 15** | 56 |  |  |  |  |
|  | R Frontal Pole |  | 3.45 | 10 | 62 | 10 |
|  | **Cluster 16** | 56 |  |  |  |  |
|  | L Postcentral Gyrus |  | 3.97 | -34 | -26 | 52 |
|  | **Cluster 17** | 48 |  |  |  |  |
|  | L Frontal Medial Cortex |  | 2.25 | -6 | 40 | -22 |
|  | **Cluster 18** | 48 |  |  |  |  |
|  | L Frontal Pole |  | 3.18 | -28 | 62 | -10 |
|  | **Cluster 19** | 48 |  |  |  |  |
|  | L Middle Temporal Gyrus, posterior division |  | 2.84 | -68 | -38 | 2 |
|  | **Cluster 20** | 40 |  |  |  |  |
|  | L Brain Stem |  | 3.26 | -14 | -26 | -38 |
|  | **Cluster 21** | 40 |  |  |  |  |
|  | R Subcallosal Cortex |  | 3.06 | 0 | 24 | -4 |
|  | **Cluster 22** | 40 |  |  |  |  |
|  | L Planum Temporale |  | 3.42 | -52 | -24 | 4 |
