## Extended Data Table 5-2 for "Neuroticism/negative emotionality is associated with increased reactivity to uncertain threat in the bed nucleus of the stria terminalis, not the amygdala" (from another version of this preprint: version 5, 2024-04-21)

**Extended Data Table 5-2.** Descriptive statistics for clusters and local extrema showing greater activation during the anticipation of Uncertain Threat relative to the implicit baseline (FDR *q*<0.05, whole-brain corrected; spatially unsmoothed data).

|  | | **mm^3^** | ***t*** | **x** | **y** | **z** |
| --- | --- | --- | --- | --- | --- | --- |
|  | **Cluster 1** | 360,632 |  |  |  |  |
|  | L Frontal Operculum Cortex |  | 12.24 | -32 | 20 | 10 |
|  | L Frontal Orbital Cortex |  | 11.50 | -30 | 28 | 0 |
|  | L Cingulate Gyrus, anterior division |  | 9.86 | -2 | 22 | 34 |
|  | L Central Opercular Cortex |  | 9.53 | -42 | 6 | 2 |
|  | L Paracingulate Gyrus |  | 8.61 | -8 | 22 | 32 |
|  | L Supramarginal Gyrus, posterior division |  | 8.59 | -56 | -50 | 36 |
|  | L Superior Frontal Gyrus |  | 8.58 | -12 | -6 | 68 |
|  | L Juxtapositional Lobule Cortex |  | 8.35 | -2 | 6 | 46 |
|  | L Angular Gyrus |  | 8.33 | -60 | -54 | 36 |
|  | L Putamen |  | 8.06 | -24 | 6 | -4 |
|  | L Brain Stem |  | 7.94 | -2 | -28 | -2 |
|  | L Frontal Pole |  | 7.58 | -36 | 48 | 30 |
|  | L Cingulate Gyrus, posterior division |  | 7.50 | -12 | -24 | 38 |
|  | L Precentral Gyrus |  | 7.37 | -32 | -6 | 50 |
|  | L Occipital Fusiform Gyrus |  | 7.37 | -14 | -90 | -12 |
|  | L Bed Nucleus of the Stria Terminalis/Caudate |  | 7.35 | -8 | 2 | 6 |
|  | L Inferior Frontal Gyrus, pars opercularis |  | 7.27 | -56 | 14 | 0 |
|  | L Parietal Operculum Cortex |  | 7.21 | -62 | -30 | 20 |
|  | L Temporal Occipital Fusiform Cortex |  | 7.20 | -34 | -56 | -22 |
|  | L Inferior Frontal Gyrus, pars triangularis |  | 7.18 | -54 | 22 | 4 |
|  | L Occipital Pole |  | 7.01 | -22 | -98 | 14 |
|  | L Supramarginal Gyrus, anterior division |  | 7.00 | -56 | -38 | 32 |
|  | L Precuneus Cortex |  | 6.81 | -10 | -70 | 38 |
|  | L Lingual Gyrus |  | 6.76 | -8 | -72 | -12 |
|  | L Postcentral Gyrus |  | 6.54 | -66 | -22 | 22 |
|  | L Middle Frontal Gyrus |  | 6.49 | -34 | 34 | 38 |
|  | L Superior Parietal Lobule |  | 6.30 | -18 | -52 | 64 |
|  | L Thalamus |  | 6.23 | -8 | -4 | 12 |
|  | L Lateral Occipital Cortex, superior division |  | 6.12 | -38 | -58 | 58 |
|  | L Bed Nucleus of the Stria Terminalis/Anterior Commissure |  | 5.99 | -8 | 2 | -6 |
|  | L Lateral Occipital Cortex, inferior division |  | 5.42 | -42 | -84 | -10 |
|  | L Insular Cortex |  | 4.49 | -38 | 2 | -2 |
|  | L Temporal Pole |  | 4.26 | -58 | 6 | -8 |
|  | L Temporal Fusiform Cortex, posterior division |  | 4.20 | -40 | -42 | -18 |
|  | L Inferior Temporal Gyrus, temporooccipital part |  | 4.11 | -46 | -62 | -10 |
|  | L Pallidum |  | 4.05 | -26 | -16 | -2 |
|  | L Bed Nucleus of the Stria Terminalis |  | 3.95 | -6 | 2 | 0 |
|  | R Frontal Operculum Cortex |  | 11.68 | 34 | 24 | 10 |
|  | R Paracingulate Gyrus |  | 11.60 | 10 | 20 | 34 |
|  | R Precentral Gyrus |  | 11.14 | 40 | -2 | 46 |
|  | R Thalamus |  | 10.72 | 10 | 0 | 10 |
|  | R Cingulate Gyrus, anterior division |  | 10.68 | 10 | 12 | 38 |
|  | R Juxtapositional Lobule Cortex |  | 10.45 | 10 | 4 | 46 |
|  | R Frontal Orbital Cortex |  | 10.30 | 32 | 22 | -8 |
|  | R Putamen |  | 10.12 | 32 | -2 | -6 |
|  | R Central Opercular Cortex |  | 10.02 | 46 | 8 | 0 |
|  | R Parietal Operculum Cortex |  | 9.85 | 52 | -30 | 28 |
|  | R Brain Stem |  | 9.84 | 4 | -28 | -2 |
|  | R Lateral Occipital Cortex, inferior division |  | 9.61 | 42 | -86 | -8 |
|  | R Middle Frontal Gyrus |  | 9.37 | 46 | 2 | 56 |
|  | R Angular Gyrus |  | 9.25 | 64 | -46 | 28 |
|  | R Supramarginal Gyrus, posterior division |  | 9.23 | 62 | -46 | 34 |
|  | R Superior Parietal Lobule |  | 9.02 | 22 | -46 | 64 |
|  | R Supramarginal Gyrus, anterior division |  | 8.94 | 54 | -30 | 32 |
|  | R Occipital Pole |  | 8.92 | 26 | -94 | 12 |
|  | R Superior Frontal Gyrus |  | 8.88 | 18 | -8 | 70 |
|  | R Temporal Pole |  | 8.87 | 50 | 10 | -4 |
|  | R Occipital Fusiform Gyrus |  | 8.62 | 38 | -68 | -12 |
|  | R Superior Temporal Gyrus, posterior division |  | 8.19 | 48 | -24 | -4 |
|  | R Inferior Frontal Gyrus, pars triangularis |  | 8.10 | 52 | 22 | 2 |
|  | R Cingulate Gyrus, posterior division |  | 7.79 | 10 | -20 | 42 |
|  | R Caudate |  | 7.60 | 12 | -2 | 16 |
|  | R Inferior Frontal Gyrus, pars opercularis |  | 7.44 | 54 | 10 | 6 |
|  | R Postcentral Gyrus |  | 7.40 | 34 | -38 | 62 |
|  | R Frontal Pole |  | 7.15 | 26 | 48 | 24 |
|  | R Bed Nucleus of the Stria Terminalis |  | 6.87 | 8 | 2 | 4 |
|  | R Middle Temporal Gyrus, posterior division |  | 6.61 | 54 | -30 | -4 |
|  | R Inferior Temporal Gyrus, temporooccipital part |  | 6.50 | 46 | -50 | -14 |
|  | R Insular Cortex |  | 6.49 | 38 | 4 | 0 |
|  | R Middle Temporal Gyrus, temporooccipital part |  | 5.93 | 46 | -56 | 6 |
|  | R Lingual Gyrus |  | 5.68 | 10 | -80 | -4 |
|  | R Lateral Occipital Cortex, superior division |  | 5.65 | 34 | -66 | 28 |
|  | R Temporal Occipital Fusiform Cortex |  | 5.63 | 38 | -48 | -18 |
|  | R Accumbens |  | 5.11 | 10 | 6 | -6 |
|  | R Pallidum |  | 5.08 | 18 | 2 | 2 |
|  | R Precuneus Cortex |  | 4.92 | 4 | -48 | 54 |
|  | R Planum Polare |  | 4.66 | 46 | -8 | -6 |
|  | R Thalamus |  | 4.35 | 0 | -10 | 0 |
|  | R Temporal Fusiform Cortex, posterior division |  | 3.80 | 32 | -34 | -24 |
|  | R Hippocampus |  | 3.79 | 34 | -30 | -6 |
|  | R Amygdala |  | 3.66 | 18 | -2 | -14 |
|  | R Heschls Gyrus (includes H1 and H2) |  | 3.08 | 48 | -22 | 8 |
|  | **Cluster 2** | 2,352 |  |  |  |  |
|  | L Middle Temporal Gyrus, temporooccipital part |  | 4.52 | -62 | -48 | 6 |
|  | L Supramarginal Gyrus, posterior division |  | 4.23 | -54 | -48 | 14 |
|  | L Middle Temporal Gyrus, posterior division |  | 3.97 | -60 | -26 | -4 |
|  | L Angular Gyrus |  | 3.19 | -64 | -50 | 10 |
|  | **Cluster 3** | 1,432 |  |  |  |  |
|  | R Precuneus Cortex |  | 5.83 | 16 | -64 | 36 |
|  | R Cuneal Cortex |  | 2.71 | 12 | -78 | 38 |
|  | **Cluster 4** | 920 |  |  |  |  |
|  | R Temporal Pole |  | 4.24 | 42 | 8 | -38 |
|  | R Middle Temporal Gyrus, anterior division |  | 3.99 | 48 | 2 | -32 |
|  | **Cluster 5** | 448 |  |  |  |  |
|  | L Inferior Temporal Gyrus, anterior division |  | 3.76 | -46 | -6 | -32 |
|  | L Temporal Pole |  | 3.69 | -38 | 4 | -36 |
|  | L Middle Temporal Gyrus, anterior division |  | 3.67 | -50 | 0 | -32 |
|  | L Middle Temporal Gyrus, posterior division |  | 3.24 | -58 | -12 | -28 |
|  | **Cluster 6** | 256 |  |  |  |  |
|  | L Frontal Pole |  | 4.73 | -32 | 54 | -12 |
|  | **Cluster 7** | 248 |  |  |  |  |
|  | L Planum Polare |  | 4.80 | -40 | -14 | -8 |
|  | **Cluster 8** | 224 |  |  |  |  |
|  | R Right Hippocampus |  | 4.60 | 30 | -38 | 2 |
|  | **Cluster 9** | 176 |  |  |  |  |
|  | L Brain Stem |  | 3.86 | -6 | -46 | -52 |
|  | **Cluster 10** | 136 |  |  |  |  |
|  | R Frontal Pole |  | 3.93 | 40 | 46 | 6 |
|  | **Cluster 11** | 128 |  |  |  |  |
|  | L Thalamus |  | 3.45 | -22 | -34 | -2 |
|  | L Hippocampus |  | 2.95 | -28 | -36 | -2 |
|  | **Cluster 12** | 120 |  |  |  |  |
|  | L Precentral Gyrus |  | 3.82 | -36 | -16 | 42 |
|  | L Postcentral Gyrus |  | 3.51 | -38 | -18 | 40 |
|  | **Cluster 13** | 112 |  |  |  |  |
|  | L Frontal Pole |  | 4.01 | -16 | 52 | -16 |
|  | **Cluster 14** | 104 |  |  |  |  |
|  | L Heschls Gyrus (includes H1 and H2) |  | 4.35 | -38 | -28 | 12 |
|  | **Cluster 15** | 96 |  |  |  |  |
|  | L Middle Frontal Gyrus |  | 4.42 | -46 | 14 | 30 |
|  | **Cluster 16** | 88 |  |  |  |  |
|  | L Middle Temporal Gyrus, posterior division |  | 3.06 | -56 | -34 | -10 |
|  | **Cluster 17** | 64 |  |  |  |  |
|  | L Middle Temporal Gyrus, posterior division |  | 2.96 | -68 | -42 | -2 |
|  | **Cluster 18** | 56 |  |  |  |  |
|  | L Temporal Pole |  | 3.13 | -46 | 8 | -34 |
|  | **Cluster 19** | 56 |  |  |  |  |
|  | L Paracingulate Gyrus |  | 2.99 | -12 | 50 | 6 |
|  | **Cluster 20** | 56 |  |  |  |  |
|  | L Inferior Frontal Gyrus, pars opercularis |  | 2.83 | -38 | 16 | 26 |
|  | L Middle Frontal Gyrus |  | 2.48 | -36 | 14 | 28 |
|  | **Cluster 21** | 48 |  |  |  |  |
|  | L Inferior Temporal Gyrus, posterior division |  | 3.65 | -56 | -20 | -30 |
|  | **Cluster 22** | 48 |  |  |  |  |
|  | L Hippocampus |  | 3.60 | -22 | -34 | -8 |
|  | **Cluster 23** | 40 |  |  |  |  |
|  | L Hippocampus |  | 3.76 | -28 | -18 | -12 |
|  | **Cluster 24** | 40 |  |  |  |  |
|  | L Precuneus Cortex |  | 2.80 | -8 | -42 | 44 |

Note: Suprathreshold activation was also evident in the right dorsal amygdala.
