## Extended Data Table 5-3 for "Neuroticism/negative emotionality is associated with increased reactivity to uncertain threat in the bed nucleus of the stria terminalis, not the amygdala" (from another version of this preprint: version 5, 2024-04-21)

**Extended Data Table 5-3.** Descriptive statistics for clusters and local extrema showing greater activation during the anticipation of Certain Threat relative to the implicit baseline (FDR *q*<0.05, whole-brain corrected; spatially unsmoothed data).

|  | | **mm^3^** | ***t*** | **x** | **y** | **z** |
| --- | --- | --- | --- | --- | --- | --- |
|  | **Cluster 1** | 67,208 |  |  |  |  |
|  | L Cingulate Gyrus, anterior division |  | 8.15 | -8 | 16 | 34 |
|  | L Superior Frontal Gyrus |  | 7.25 | -14 | -2 | 68 |
|  | L Paracingulate Gyrus |  | 6.75 | -4 | 12 | 42 |
|  | L Juxtapositional Lobule Cortex |  | 6.25 | -6 | 4 | 56 |
|  | L Precentral Gyrus |  | 5.97 | -28 | -8 | 46 |
|  | L Middle Frontal Gyrus |  | 4.88 | -40 | 0 | 50 |
|  | L Postcentral Gyrus |  | 3.49 | -40 | -18 | 40 |
|  | R Cingulate Gyrus, anterior division |  | 8.37 | 10 | 12 | 38 |
|  | R Paracingulate Gyrus |  | 7.52 | 12 | 20 | 34 |
|  | R Superior Frontal Gyrus |  | 7.07 | 14 | -6 | 68 |
|  | R Juxtapositional Lobule Cortex |  | 6.68 | 8 | 6 | 58 |
|  | R Precentral Gyrus |  | 6.45 | 48 | 2 | 46 |
|  | R Supramarginal Gyrus, posterior division |  | 6.14 | 64 | -40 | 26 |
|  | R Lateral Occipital Cortex, superior division |  | 6.00 | 18 | -76 | 42 |
|  | R Superior Temporal Gyrus, posterior division |  | 5.87 | 48 | -24 | -4 |
|  | R Postcentral Gyrus |  | 5.82 | 44 | -28 | 42 |
|  | R Superior Parietal Lobule |  | 5.78 | 34 | -42 | 58 |
|  | R Cingulate Gyrus, posterior division |  | 5.68 | 14 | -28 | 40 |
|  | R Angular Gyrus |  | 5.46 | 60 | -46 | 32 |
|  | R Precuneus Cortex |  | 5.34 | 14 | -78 | 42 |
|  | R Middle Temporal Gyrus, temporooccipital part |  | 5.26 | 50 | -42 | 8 |
|  | R Middle Frontal Gyrus |  | 5.20 | 42 | 0 | 54 |
|  | R Parietal Operculum Cortex |  | 5.15 | 56 | -28 | 24 |
|  | R Middle Temporal Gyrus, posterior division |  | 4.66 | 48 | -22 | -8 |
|  | R Supramarginal Gyrus, anterior division |  | 4.59 | 58 | -24 | 32 |
|  | R Occipital Pole |  | 4.31 | 18 | -90 | 34 |
|  | R Cuneal Cortex |  | 4.13 | 12 | -80 | 38 |
|  | R Planum Temporale |  | 3.36 | 62 | -32 | 20 |
|  | **Cluster 2** | 41,944 |  |  |  |  |
|  | L Putamen |  | 6.71 | -20 | 12 | -2 |
|  | L Thalamus |  | 5.98 | -2 | -2 | 4 |
|  | L Lingual Gyrus |  | 5.83 | -10 | -72 | -10 |
|  | L Bed Nucleus of the Stria Terminalis |  | 5.81 | -6 | 2 | 0 |
|  | L Pallidum |  | 5.68 | -20 | 2 | -4 |
|  | L Occipital Fusiform Gyrus |  | 5.57 | -30 | -68 | -18 |
|  | L Temporal Occipital Fusiform Cortex |  | 5.54 | -26 | -60 | -16 |
|  | L Caudate |  | 5.35 | -8 | 0 | 8 |
|  | L Brain Stem |  | 5.18 | -2 | -28 | -2 |
|  | L Insular Cortex |  | 2.95 | -32 | 6 | 8 |
|  | L Accumbens |  | 2.83 | -6 | 12 | -6 |
|  | R Temporal Occipital Fusiform Cortex |  | 5.86 | 24 | -54 | -16 |
|  | R Thalamus |  | 5.83 | 2 | -16 | -2 |
|  | R Lingual Gyrus |  | 5.46 | 12 | -70 | -12 |
|  | R Brain Stem |  | 5.24 | 4 | -28 | -2 |
|  | R Occipital Fusiform Gyrus |  | 4.71 | 20 | -68 | -14 |
|  | R Thalamus |  | 3.35 | 0 | -10 | 2 |
|  | **Cluster 3** | 20,704 |  |  |  |  |
|  | L Supramarginal Gyrus, posterior division |  | 6.43 | -56 | -50 | 40 |
|  | L Precuneus Cortex |  | 5.43 | -10 | -70 | 36 |
|  | L Supramarginal Gyrus, anterior division |  | 5.29 | -58 | -34 | 40 |
|  | L Lateral Occipital Cortex, superior division |  | 5.08 | -10 | -64 | 62 |
|  | L Angular Gyrus |  | 5.07 | -60 | -54 | 36 |
|  | L Cuneal Cortex |  | 4.96 | -12 | -80 | 38 |
|  | L Parietal Operculum Cortex |  | 4.88 | -58 | -36 | 22 |
|  | L Superior Parietal Lobule |  | 4.80 | -36 | -48 | 48 |
|  | L Postcentral Gyrus |  | 4.68 | -40 | -32 | 48 |
|  | L Superior Temporal Gyrus, posterior division |  | 2.77 | -66 | -34 | 14 |
|  | **Cluster 4** | 15,232 |  |  |  |  |
|  | R Putamen |  | 7.94 | 24 | 10 | -4 |
|  | R Caudate |  | 7.33 | 10 | 2 | 10 |
|  | R Pallidum |  | 7.19 | 20 | 4 | 0 |
|  | R Bed Nucleus of the Stria Terminalis/Caudate |  | 6.50 | 8 | 6 | 0 |
|  | R Bed Nucleus of the Stria Terminalis |  | 6.36 | 8 | 2 | 4 |
|  | R Central Opercular Cortex |  | 6.17 | 48 | 8 | -2 |
|  | R Frontal Operculum Cortex |  | 6.07 | 34 | 24 | 8 |
|  | R Precentral Gyrus |  | 5.61 | 62 | 4 | 12 |
|  | R Thalamus |  | 5.56 | 10 | -2 | 12 |
|  | R Insular Cortex |  | 5.47 | 34 | 16 | 8 |
|  | R Frontal Orbital Cortex |  | 5.25 | 32 | 30 | -2 |
|  | R Temporal Pole |  | 4.38 | 54 | 10 | -2 |
|  | R Accumbens |  | 4.34 | 10 | 12 | -6 |
|  | R Frontal Pole |  | 4.32 | 50 | 34 | -4 |
|  | R Inferior Frontal Gyrus, pars triangularis |  | 4.17 | 54 | 26 | 6 |
|  | R Inferior Frontal Gyrus, pars opercularis |  | 3.66 | 56 | 10 | 6 |
|  | **Cluster 5** | 7,880 |  |  |  |  |
|  | L Frontal Pole |  | 6.03 | -30 | 52 | 28 |
|  | L Middle Frontal Gyrus |  | 5.73 | -34 | 34 | 38 |
|  | **Cluster 6** | 5,616 |  |  |  |  |
|  | R Frontal Pole |  | 6.39 | 34 | 44 | 28 |
|  | R Middle Frontal Gyrus |  | 6.35 | 30 | 32 | 34 |
|  | **Cluster 7** | 5,536 |  |  |  |  |
|  | L Frontal Operculum Cortex |  | 7.52 | -36 | 12 | 6 |
|  | L Central Opercular Cortex |  | 5.93 | -42 | 8 | 2 |
|  | L Insular Cortex |  | 5.57 | -30 | 26 | 6 |
|  | L Inferior Frontal Gyrus, pars opercularis |  | 5.03 | -52 | 14 | 2 |
|  | L Frontal Orbital Cortex |  | 4.65 | -30 | 28 | 0 |
|  | L Temporal Pole |  | 4.21 | -52 | 12 | -6 |
|  | L Precentral Gyrus |  | 3.35 | -52 | 8 | 4 |
|  | L Inferior Frontal Gyrus, pars triangularis |  | 2.82 | -52 | 24 | -4 |
|  | **Cluster 8** | 1,224 |  |  |  |  |
|  | L Cingulate Gyrus, posterior division |  | 4.52 | -14 | -28 | 38 |
|  | L Precuneus Cortex |  | 4.22 | -10 | -44 | 50 |
|  | L Precentral Gyrus |  | 4.20 | -16 | -32 | 42 |
|  | **Cluster 9** | 856 |  |  |  |  |
|  | R Thalamus |  | 4.39 | 16 | -18 | 6 |
|  | **Cluster 10** | 832 |  |  |  |  |
|  | R Lateral Occipital Cortex, inferior division |  | 4.57 | 44 | -62 | -12 |
|  | R Inferior Temporal Gyrus, temporooccipital part |  | 3.97 | 52 | -60 | -12 |
|  | R Middle Temporal Gyrus, temporooccipital part |  | 3.22 | 48 | -56 | 2 |
|  | **Cluster 11** | 664 |  |  |  |  |
|  | R Putamen |  | 5.23 | 30 | -18 | 2 |
|  | R Amygdala (basolateral nucleus) |  | 5.11 | 30 | -8 | -14 |
|  | R Hippocampus |  | 3.34 | 26 | -16 | -12 |
|  | **Cluster 12** | 424 |  |  |  |  |
|  | R Occipital Pole |  | 4.07 | 24 | -92 | 10 |
|  | R Lateral Occipital Cortex, superior division |  | 3.30 | 30 | -84 | 18 |
|  | **Cluster 13** | 376 |  |  |  |  |
|  | L Cingulate Gyrus, posterior division |  | 3.69 | -2 | -28 | 26 |
|  | R Cingulate Gyrus, posterior division |  | 3.83 | 8 | -26 | 28 |
|  | **Cluster 14** | 360 |  |  |  |  |
|  | L Lingual Gyrus |  | 4.50 | -10 | -84 | -8 |
|  | L Occipital Fusiform Gyrus |  | 3.42 | -16 | -78 | -10 |
|  | **Cluster 15** | 304 |  |  |  |  |
|  | L Putamen |  | 5.68 | -32 | -12 | -10 |
|  | L Amygdala (central nucleus) |  | 4.14 | -24 | -12 | -10 |
|  | L Hippocampus |  | 3.58 | -30 | -16 | -14 |
|  | **Cluster 16** | 296 |  |  |  |  |
|  | L Hippocampus |  | 4.17 | -20 | -42 | 2 |
|  | L Thalamus |  | 3.83 | -22 | -36 | 0 |
|  | **Cluster 17** | 296 |  |  |  |  |
|  | L Postcentral Gyrus |  | 3.53 | -66 | -22 | 22 |
|  | **Cluster 18** | 264 |  |  |  |  |
|  | R Hippocampus |  | 3.98 | 24 | -36 | 4 |
|  | R Thalamus |  | 3.75 | 20 | -34 | 6 |
|  | **Cluster 19** | 184 |  |  |  |  |
|  | L Frontal Pole |  | 4.18 | -30 | 50 | -14 |
|  | **Cluster 20** | 176 |  |  |  |  |
|  | L Brain Stem |  | 3.74 | -6 | -38 | -44 |
|  | **Cluster 21** | 168 |  |  |  |  |
|  | R Brain Stem |  | 3.87 | 12 | -26 | -14 |
|  | **Cluster 22** | 168 |  |  |  |  |
|  | L Lateral Occipital Cortex, inferior division |  | 4.32 | -42 | -80 | -4 |
|  | **Cluster 23** | 160 |  |  |  |  |
|  | R Cingulate Gyrus, anterior division |  | 3.93 | 4 | -8 | 40 |
|  | **Cluster 24** | 144 |  |  |  |  |
|  | L Postcentral Gyrus |  | 3.85 | -56 | -18 | 30 |
|  | **Cluster 25** | 136 |  |  |  |  |
|  | R Superior Frontal Gyrus |  | 4.34 | 4 | 56 | 34 |
|  | **Cluster 26** | 128 |  |  |  |  |
|  | R Hippocampus |  | 4.83 | 34 | -34 | -6 |
|  | **Cluster 27** | 128 |  |  |  |  |
|  | L Supramarginal Gyrus, posterior division |  | 3.60 | -52 | -48 | 12 |
|  | **Cluster 28** | 128 |  |  |  |  |
|  | R Inferior Frontal Gyrus, pars opercularis |  | 3.33 | 38 | 12 | 26 |
|  | **Cluster 29** | 120 |  |  |  |  |
|  | L Paracingulate Gyrus |  | 3.64 | -10 | 50 | 8 |
|  | **Cluster 30** | 112 |  |  |  |  |
|  | R Lateral Occipital Cortex, inferior division |  | 3.84 | 40 | -80 | -4 |
|  | **Cluster 31** | 104 |  |  |  |  |
|  | R Brain Stem |  | 3.62 | 8 | -38 | -48 |
|  | **Cluster 32** | 104 |  |  |  |  |
|  | R Lateral Occipital Cortex, inferior division |  | 2.99 | 50 | -66 | 6 |
|  | **Cluster 33** | 96 |  |  |  |  |
|  | R Lateral Occipital Cortex, inferior division |  | 3.78 | 42 | -68 | 10 |
|  | R Lateral Occipital Cortex, superior division |  | 3.60 | 40 | -66 | 22 |
|  | **Cluster 34** | 88 |  |  |  |  |
|  | R Temporal Pole |  | 4.02 | 36 | 8 | -36 |
|  | **Cluster 35** | 88 |  |  |  |  |
|  | L Heschls Gyrus (includes H1 and H2) |  | 4.22 | -38 | -28 | 12 |
|  | **Cluster 36** | 88 |  |  |  |  |
|  | L Central Opercular Cortex |  | 3.38 | -54 | -20 | 18 |
|  | **Cluster 37** | 88 |  |  |  |  |
|  | R Postcentral Gyrus |  | 3.27 | 22 | -36 | 70 |
|  | **Cluster 38** | 88 |  |  |  |  |
|  | L Superior Frontal Gyrus |  | 3.57 | -4 | 20 | 64 |
|  | **Cluster 39** | 80 |  |  |  |  |
|  | L Middle Temporal Gyrus, temporooccipital part |  | 3.49 | -66 | -50 | 6 |
|  | **Cluster 40** | 80 |  |  |  |  |
|  | L Lateral Occipital Cortex, inferior division |  | 3.25 | -40 | -70 | 14 |
|  | **Cluster 41** | 80 |  |  |  |  |
|  | L Cingulate Gyrus, posterior division |  | 3.62 | -4 | -18 | 28 |
|  | R Cingulate Gyrus, posterior division |  | 3.38 | 2 | -22 | 30 |
|  | **Cluster 42** | 80 |  |  |  |  |
|  | L Superior Frontal Gyrus |  | 3.85 | -12 | 32 | 52 |
|  | **Cluster 43** | 72 |  |  |  |  |
|  | L Hippocampus |  | 3.64 | -34 | -30 | -8 |
|  | **Cluster 44** | 72 |  |  |  |  |
|  | L Middle Temporal Gyrus, temporooccipital part |  | 2.95 | -58 | -50 | -4 |
|  | **Cluster 45** | 72 |  |  |  |  |
|  | R Frontal Pole |  | 3.24 | 42 | 50 | 2 |
|  | **Cluster 46** | 72 |  |  |  |  |
|  | L Precentral Gyrus |  | 5.19 | -56 | 0 | 14 |
|  | **Cluster 47** | 64 |  |  |  |  |
|  | L Putamen |  | 4.08 | -32 | -20 | -4 |
|  | **Cluster 48** | 64 |  |  |  |  |
|  | L Lingual Gyrus |  | 3.68 | -22 | -50 | -2 |
|  | **Cluster 49** | 64 |  |  |  |  |
|  | R Lateral Occipital Cortex, inferior division |  | 3.28 | 58 | -62 | 6 |
|  | **Cluster 50** | 64 |  |  |  |  |
|  | L Occipital Pole |  | 3.25 | -20 | -94 | 10 |
|  | **Cluster 51** | 64 |  |  |  |  |
|  | L Central Opercular Cortex |  | 4.40 | -44 | 0 | 12 |
|  | **Cluster 52** | 64 |  |  |  |  |
|  | L Precentral Gyrus |  | 3.65 | -60 | 2 | 20 |
|  | **Cluster 53** | 64 |  |  |  |  |
|  | R Postcentral Gyrus |  | 3.50 | 52 | -16 | 32 |
|  | **Cluster 54** | 56 |  |  |  |  |
|  | R Brain Stem |  | 3.43 | 2 | -34 | -48 |
|  | **Cluster 55** | 56 |  |  |  |  |
|  | L Brain Stem |  | 3.39 | -10 | -38 | -30 |
|  | **Cluster 56** | 56 |  |  |  |  |
|  | R Brain Stem |  | 3.27 | 4 | -26 | -22 |
|  | **Cluster 57** | 56 |  |  |  |  |
|  | L Lateral Occipital Cortex, inferior division |  | 3.40 | -48 | -74 | -10 |
|  | **Cluster 58** | 56 |  |  |  |  |
|  | L Hippocampus |  | 3.74 | -34 | -22 | -10 |
|  | **Cluster 59** | 56 |  |  |  |  |
|  | L Frontal Pole |  | 3.79 | -32 | 38 | -12 |
|  | **Cluster 60** | 56 |  |  |  |  |
|  | R Middle Temporal Gyrus, temporooccipital part |  | 3.10 | 56 | -56 | 6 |
|  | **Cluster 61** | 56 |  |  |  |  |
|  | L Frontal Pole |  | 3.88 | -38 | 60 | 6 |
|  | **Cluster 62** | 56 |  |  |  |  |
|  | L Lateral Occipital Cortex, inferior division |  | 2.96 | -56 | -66 | 8 |
|  | **Cluster 63** | 56 |  |  |  |  |
|  | R Occipital Pole |  | 3.61 | 20 | -98 | 10 |
|  | **Cluster 64** | 56 |  |  |  |  |
|  | L Angular Gyrus |  | 3.11 | -46 | -56 | 14 |
|  | **Cluster 65** | 56 |  |  |  |  |
|  | R Middle Temporal Gyrus, temporooccipital part |  | 3.48 | 54 | -52 | 10 |
|  | **Cluster 66** | 56 |  |  |  |  |
|  | R Cingulate Gyrus, anterior division |  | 3.52 | 6 | 28 | 18 |
|  | **Cluster 67** | 56 |  |  |  |  |
|  | L Cingulate Gyrus, anterior division |  | 3.85 | -2 | -6 | 38 |
|  | **Cluster 68** | 56 |  |  |  |  |
|  | R Precentral Gyrus |  | 3.36 | 24 | -18 | 74 |
|  | **Cluster 69** | 48 |  |  |  |  |
|  | L Inferior Temporal Gyrus, temporooccipital part |  | 3.45 | -44 | -48 | -12 |
|  | **Cluster 70** | 48 |  |  |  |  |
|  | R Middle Temporal Gyrus, posterior division |  | 3.59 | 68 | -20 | -4 |
|  | **Cluster 71** | 48 |  |  |  |  |
|  | L Lateral Occipital Cortex, inferior division |  | 3.65 | -48 | -66 | 2 |
|  | **Cluster 72** | 48 |  |  |  |  |
|  | R Lateral Occipital Cortex, inferior division |  | 3.69 | 40 | -88 | 2 |
|  | **Cluster 73** | 48 |  |  |  |  |
|  | L Supramarginal Gyrus, posterior division |  | 2.93 | -64 | -46 | 8 |
|  | **Cluster 74** | 48 |  |  |  |  |
|  | L Central Opercular Cortex |  | 2.88 | -34 | 4 | 14 |
|  | **Cluster 75** | 48 |  |  |  |  |
|  | R Inferior Frontal Gyrus, pars opercularis |  | 3.21 | 48 | 14 | 20 |
|  | **Cluster 76** | 48 |  |  |  |  |
|  | R Inferior Frontal Gyrus, pars opercularis |  | 4.38 | 56 | 16 | 20 |
|  | R Precentral Gyrus |  | 2.80 | 56 | 10 | 20 |
|  | **Cluster 77** | 48 |  |  |  |  |
|  | R Superior Frontal Gyrus |  | 4.36 | 12 | 28 | 60 |
|  | **Cluster 78** | 40 |  |  |  |  |
|  | R Brain Stem |  | 4.06 | 8 | -46 | -54 |
|  | **Cluster 79** | 40 |  |  |  |  |
|  | L Brain Stem |  | 3.36 | -2 | -32 | -24 |
|  | **Cluster 80** | 40 |  |  |  |  |
|  | L Middle Temporal Gyrus, posterior division |  | 3.29 | -62 | -34 | -18 |
|  | **Cluster 81** | 40 |  |  |  |  |
|  | L Lateral Occipital Cortex, inferior division |  | 3.13 | -38 | -72 | -4 |
|  | **Cluster 82** | 40 |  |  |  |  |
|  | R Paracingulate Gyrus |  | 3.56 | 14 | 52 | 6 |
|  | **Cluster 83** | 40 |  |  |  |  |
|  | R Middle Temporal Gyrus, temporooccipital part |  | 3.16 | 62 | -54 | 12 |
|  | **Cluster 84** | 40 |  |  |  |  |
|  | R Thalamus |  | 3.73 | 8 | -20 | 16 |
|  | **Cluster 85** | 40 |  |  |  |  |
|  | L Caudate |  | 3.67 | -18 | -16 | 24 |
|  | **Cluster 86** | 40 |  |  |  |  |
|  | R Frontal Pole |  | 2.80 | 14 | 58 | 28 |
|  | **Cluster 87** | 40 |  |  |  |  |
|  | R Angular Gyrus |  | 3.20 | 52 | -52 | 46 |

Note: Suprathreshold activation was also evident in the L Dorsal Amygdala in the region of the Ce.
