## Extended Data Table 5-4 for "Neuroticism/negative emotionality is associated with increased reactivity to uncertain threat in the bed nucleus of the stria terminalis, not the amygdala" (from another version of this preprint: version 5, 2024-04-21)

**Extended Data Table 5-4.** Descriptive statistics for clusters and local extrema showing greater activation during the presentation of ‘threat-related’ faces relative to the implicit baseline (FDR *q*<0.05, whole-brain corrected; spatially unsmoothed data).

|  | | **mm^3^** | ***t*** | **x** | **y** | **z** |
| --- | --- | --- | --- | --- | --- | --- |
|  | **Cluster 1** | 648,072 |  |  |  |  |
|  | L Postcentral Gyrus |  | 15.78 | -34 | -28 | 60 |
|  | L Amygdala (cortical, amygdalohippocampal area) |  | 14.14 | -16 | -4 | -16 |
|  | L Precentral Gyrus |  | 13.73 | -12 | -32 | 68 |
|  | L Superior Parietal Lobule |  | 12.92 | -36 | -42 | 64 |
|  | L Intracalcarine Cortex |  | 12.78 | -6 | -72 | 12 |
|  | L Superior Frontal Gyrus |  | 12.55 | -6 | -8 | 74 |
|  | L Supramarginal Gyrus, anterior division |  | 11.40 | -54 | -30 | 52 |
|  | L Temporal Occipital Fusiform Cortex |  | 11.40 | -44 | -48 | -20 |
|  | L Lateral Occipital Cortex, inferior division |  | 11.18 | -56 | -70 | 10 |
|  | L Middle Frontal Gyrus |  | 10.76 | -44 | 0 | 58 |
|  | L Juxtapositional Lobule Cortex |  | 10.56 | -2 | -8 | 66 |
|  | L Supracalcarine Cortex |  | 10.37 | -2 | -84 | 10 |
|  | L Middle Temporal Gyrus, temporooccipital part |  | 9.39 | -58 | -60 | 8 |
|  | L Angular Gyrus |  | 9.33 | -60 | -52 | 12 |
|  | L Thalamus |  | 9.29 | -8 | -2 | 10 |
|  | L Lingual Gyrus |  | 9.10 | -8 | -78 | -14 |
|  | L Cingulate Gyrus, posterior division |  | 8.97 | -6 | -52 | 32 |
|  | L Lateral Occipital Cortex, superior division |  | 8.69 | -36 | -58 | 58 |
|  | L Supramarginal Gyrus, posterior division |  | 8.65 | -52 | -48 | 10 |
|  | L Paracingulate Gyrus |  | 8.39 | -4 | 16 | 50 |
|  | L Caudate |  | 8.20 | -12 | -6 | 18 |
|  | L Cingulate Gyrus, anterior division |  | 8.05 | -8 | 16 | 36 |
|  | L Precuneus Cortex |  | 7.83 | -6 | -56 | 56 |
|  | L Parietal Operculum Cortex |  | 7.74 | -58 | -40 | 24 |
|  | L Cuneal Cortex |  | 7.71 | -10 | -80 | 30 |
|  | L Insular Cortex |  | 7.36 | -40 | 0 | -16 |
|  | L Occipital Pole |  | 7.23 | -6 | -90 | 16 |
|  | L Pallidum |  | 7.02 | -16 | 2 | 0 |
|  | L Temporal Pole |  | 6.97 | -36 | 4 | -20 |
|  | L Central Opercular Cortex |  | 6.95 | -56 | -18 | 20 |
|  | L Frontal Orbital Cortex |  | 6.72 | -38 | 30 | -16 |
|  | L Bed Nucleus of the Stria Terminalis |  | 6.38 | -4 | 4 | -2 |
|  | L Planum Polare |  | 6.28 | -42 | -2 | -14 |
|  | L Inferior Frontal Gyrus, pars triangularis |  | 6.21 | -50 | 24 | 4 |
|  | L Inferior Frontal Gyrus, pars opercularis |  | 6.16 | -58 | 14 | 24 |
|  | L Putamen |  | 5.96 | -30 | -22 | 2 |
|  | L Frontal Medial Cortex |  | 5.94 | -2 | 54 | -12 |
|  | L Frontal Pole |  | 5.87 | -18 | 40 | 42 |
|  | L Middle Temporal Gyrus, posterior division |  | 5.76 | -66 | -32 | 0 |
|  | L Temporal Fusiform Cortex, posterior division |  | 5.60 | -42 | -30 | -18 |
|  | L Planum Temporale |  | 5.52 | -42 | -32 | 8 |
|  | L Frontal Operculum Cortex |  | 5.30 | -42 | 22 | 4 |
|  | L Heschls Gyrus (includes H1 and H2) |  | 5.27 | -52 | -24 | 10 |
|  | L Temporal Fusiform Cortex, anterior division |  | 5.17 | -30 | 0 | -36 |
|  | L Accumbens |  | 5.05 | -8 | 16 | -6 |
|  | L Occipital Fusiform Gyrus |  | 4.89 | -40 | -70 | -16 |
|  | L Superior Temporal Gyrus, posterior division |  | 4.89 | -62 | -30 | 0 |
|  | L Hippocampus |  | 4.85 | -26 | -34 | -6 |
|  | L Brainstem |  | 4.35 | -6 | -18 | -20 |
|  | L Inferior Temporal Gyrus, posterior division |  | 4.24 | -46 | -24 | -22 |
|  | L Subcallosal Cortex |  | 4.10 | -2 | 30 | -10 |
|  | L Superior Temporal Gyrus, anterior division |  | 3.30 | -62 | 0 | -8 |
|  | L Middle Temporal Gyrus, anterior division |  | 3.00 | -62 | 0 | -22 |
|  | R Amygdala (cortical, amygdalohippocampal area) |  | 16.14 | 16 | -4 | -18 |
|  | R Intracalcarine Cortex |  | 14.97 | 10 | -72 | 14 |
|  | R Precentral Gyrus |  | 12.96 | 14 | -32 | 64 |
|  | R Postcentral Gyrus |  | 12.85 | 16 | -30 | 68 |
|  | R Supracalcarine Cortex |  | 12.40 | 2 | -74 | 12 |
|  | R Middle Frontal Gyrus |  | 12.18 | 44 | 0 | 58 |
|  | R Middle Temporal Gyrus, temporooccipital part |  | 12.10 | 48 | -58 | 6 |
|  | R Supramarginal Gyrus, posterior division |  | 12.05 | 48 | -40 | 10 |
|  | R Cingulate Gyrus, posterior division |  | 11.79 | 6 | -52 | 24 |
|  | R Superior Parietal Lobule |  | 11.39 | 26 | -42 | 66 |
|  | R Lateral Occipital Cortex, inferior division |  | 11.31 | 56 | -62 | 6 |
|  | R Central Opercular Cortex |  | 10.80 | 40 | -2 | 14 |
|  | R Superior Frontal Gyrus |  | 10.79 | 18 | -8 | 72 |
|  | R Inferior Temporal Gyrus, temporooccipital part |  | 10.68 | 46 | -42 | -20 |
|  | R Juxtapositional Lobule Cortex |  | 10.17 | 4 | 4 | 56 |
|  | R Precuneus Cortex |  | 10.11 | 2 | -62 | 34 |
|  | R Cuneal Cortex |  | 10.08 | 6 | -78 | 22 |
|  | R Lingual Gyrus |  | 10.07 | 6 | -66 | 2 |
|  | R Temporal Occipital Fusiform Cortex |  | 9.69 | 48 | -48 | -24 |
|  | R Supramarginal Gyrus, anterior division |  | 9.41 | 62 | -28 | 24 |
|  | R Thalamus |  | 9.38 | 10 | -2 | 12 |
|  | R Paracingulate Gyrus |  | 8.97 | 0 | 16 | 48 |
|  | R Caudate |  | 8.69 | 14 | -8 | 20 |
|  | R Angular Gyrus |  | 8.58 | 52 | -56 | 22 |
|  | R Thalamus |  | 8.56 | 0 | -8 | 10 |
|  | R Putamen |  | 8.21 | 30 | -18 | 0 |
|  | R Cingulate Gyrus, anterior division |  | 8.04 | 6 | -8 | 30 |
|  | R Inferior Frontal Gyrus, pars opercularis |  | 7.88 | 52 | 16 | 32 |
|  | R Middle Temporal Gyrus, posterior division |  | 7.66 | 66 | -36 | 2 |
|  | R Insular Cortex |  | 7.63 | 40 | 4 | -16 |
|  | R Parietal Operculum Cortex |  | 7.53 | 54 | -32 | 28 |
|  | R Frontal Pole |  | 7.46 | 2 | 56 | -12 |
|  | R Superior Temporal Gyrus, posterior division |  | 7.24 | 52 | -16 | -6 |
|  | R Frontal Medial Cortex |  | 6.91 | 2 | 50 | -16 |
|  | R Inferior Frontal Gyrus, pars triangularis |  | 6.88 | 54 | 26 | -6 |
|  | R Lateral Occipital Cortex, superior division |  | 6.69 | 58 | -60 | 38 |
|  | R Hippocampus |  | 6.36 | 24 | -36 | -6 |
|  | R Temporal Fusiform Cortex, anterior division |  | 6.15 | 38 | -2 | -38 |
|  | R Temporal Pole |  | 6.11 | 32 | 8 | -24 |
|  | R Pallidum |  | 6.07 | 20 | 2 | 2 |
|  | R Frontal Orbital Cortex |  | 5.91 | 48 | 22 | -12 |
|  | R Planum Temporale |  | 5.80 | 38 | -32 | 14 |
|  | R Heschls Gyrus (includes H1 and H2) |  | 5.57 | 54 | -14 | 8 |
|  | R Frontal Operculum Cortex |  | 5.48 | 42 | 26 | 0 |
|  | R Parahippocampal Gyrus, anterior division |  | 5.19 | 30 | -4 | -34 |
|  | R Bed Nucleus of the Stria Terminalis |  | 5.08 | 6 | 4 | 0 |
|  | R Inferior Temporal Gyrus, posterior division |  | 5.06 | 44 | -28 | -18 |
|  | R Middle Temporal Gyrus, anterior division |  | 4.40 | 60 | 2 | -16 |
|  | R Accumbens |  | 4.16 | 6 | 16 | -2 |
|  | R Planum Polare |  | 4.13 | 42 | -8 | -10 |
|  | R Brainstem |  | 4.00 | 4 | -42 | -22 |
|  | R Superior Temporal Gyrus, anterior division |  | 3.88 | 62 | 0 | -12 |
|  | R Subcallosal Cortex |  | 3.53 | 4 | 30 | -10 |
|  | **Cluster 2** | 552 |  |  |  |  |
|  | L Brainstem |  | 3.75 | -10 | -36 | -44 |
|  | R Brainstem |  | 3.86 | 2 | -30 | -48 |
|  | **Cluster 3** | 296 |  |  |  |  |
|  | L Brainstem |  | 3.60 | -16 | -30 | -30 |
|  | **Cluster 4** | 248 |  |  |  |  |
|  | L Middle Temporal Gyrus, anterior division |  | 4.22 | -50 | -4 | -28 |
|  | L Temporal Pole |  | 3.00 | -50 | 6 | -26 |
|  | **Cluster 5** | 240 |  |  |  |  |
|  | R Brainstem |  | 3.82 | 12 | -26 | -42 |
|  | **Cluster 6** | 224 |  |  |  |  |
|  | L Temporal Pole |  | 3.85 | -52 | 4 | -16 |
|  | **Cluster 7** | 160 |  |  |  |  |
|  | R Brainstem |  | 3.48 | 12 | -22 | -36 |
|  | **Cluster 8** | 136 |  |  |  |  |
|  | L Brainstem |  | 3.31 | -8 | -28 | -36 |
|  | R Brainstem |  | 2.68 | 2 | -32 | -36 |
|  | **Cluster 9** | 104 |  |  |  |  |
|  | L Frontal Pole |  | 4.09 | -26 | 54 | -14 |
|  | **Cluster 10** | 96 |  |  |  |  |
|  | L Temporal Pole |  | 3.94 | -44 | 12 | -28 |
|  | **Cluster 11** | 96 |  |  |  |  |
|  | R Inferior Temporal Gyrus, posterior division |  | 3.48 | 62 | -30 | -18 |
|  | R Middle Temporal Gyrus, posterior division |  | 3.18 | 64 | -32 | -16 |
|  | **Cluster 12** | 88 |  |  |  |  |
|  | L Temporal Pole |  | 3.13 | -48 | 4 | -22 |
|  | L Superior Temporal Gyrus, anterior division |  | 2.44 | -48 | 0 | -18 |
|  | **Cluster 13** | 80 |  |  |  |  |
|  | R Precentral Gyrus |  | 4.44 | 62 | 2 | 12 |
|  | **Cluster 14** | 72 |  |  |  |  |
|  | L Brainstem |  | 3.27 | -8 | -38 | -32 |
|  | **Cluster 15** | 72 |  |  |  |  |
|  | L Subcallosal Cortex |  | 2.16 | -2 | 24 | -26 |
|  | R Subcallosal Cortex |  | 3.54 | 4 | 24 | -26 |
|  | **Cluster 16** | 72 |  |  |  |  |
|  | R Frontal Pole |  | 4.48 | 22 | 42 | -16 |
|  | **Cluster 17** | 64 |  |  |  |  |
|  | R Brainstem |  | 3.45 | 6 | -36 | -48 |
|  | **Cluster 18** | 64 |  |  |  |  |
|  | R Inferior Temporal Gyrus, posterior division |  | 2.86 | 46 | -12 | -32 |
|  | **Cluster 19** | 64 |  |  |  |  |
|  | L Frontal Pole |  | 3.23 | -14 | 68 | -2 |
|  | **Cluster 20** | 64 |  |  |  |  |
|  | L Caudate |  | 2.84 | -18 | 20 | 10 |
|  | **Cluster 21** | 56 |  |  |  |  |
|  | L Frontal Pole |  | 2.80 | -42 | 52 | 0 |
|  | **Cluster 22** | 48 |  |  |  |  |
|  | R Brainstem |  | 3.46 | 0 | -18 | -38 |
|  | **Cluster 23** | 48 |  |  |  |  |
|  | L Inferior Temporal Gyrus, posterior division |  | 2.97 | -60 | -36 | -20 |
|  | **Cluster 24** | 48 |  |  |  |  |
|  | L Middle Temporal Gyrus, posterior division |  | 2.51 | -64 | -28 | -18 |
|  | L Inferior Temporal Gyrus, posterior division |  | 2.33 | -62 | -28 | -22 |
|  | **Cluster 25** | 48 |  |  |  |  |
|  | L Frontal Pole |  | 3.45 | -18 | 52 | -16 |
|  | **Cluster 26** | 40 |  |  |  |  |
|  | L Temporal Pole |  | 3.43 | -46 | 16 | -38 |
|  | **Cluster 27** | 40 |  |  |  |  |
|  | R Brainstem |  | 2.96 | 0 | -18 | -32 |
|  | **Cluster 28** | 40 |  |  |  |  |
|  | L Frontal Pole |  | 2.11 | -26 | 50 | 0 |
|  | **Cluster 29** | 40 |  |  |  |  |
|  | L Heschls Gyrus (includes H1 and H2) |  | 2.68 | -42 | -20 | 4 |

Note: Suprathreshold activation was also evident bilaterally in the dorsal amygdala.
