## Extended Data Table 7-1 for "Neuroticism/negative emotionality is associated with increased reactivity to uncertain threat in the bed nucleus of the stria terminalis, not the amygdala" (from another version of this preprint: version 5, 2024-04-21)

**Extended Data Table 7-1.** Descriptive statistics for EAc reactivity to the threat-anticipation and threat-perception paradigms (anatomically defined ROIs; spatially unsmoothed data).

| **Contrast** | ***N*** | ***M*** | ***SD*** | ***SEM*** |
| --- | --- | --- | --- | --- |
| BST, Threat minus Safety Anticipation | 220 | 0.74 | 1.17 | 0.08 |
| Ce, Threat minus Safety Anticipation | 220 | 0.40 | 0.74 | 0.05 |
| BST, Uncertain-Threat Anticipation^a^ | 220 | 0.27 | 0.73 | 0.05 |
| Ce, Uncertain-Threat Anticipation^a^ | 220 | 0.11 | 0.51 | 0.03 |
| BST, Certain-Threat Anticipation^a^ | 220 | 0.23 | 0.64 | 0.04 |
| Ce, Certain-Threat Anticipation^a^ | 220 | 0.12 | 0.48 | 0.03 |
| BST, ‘Threat-Related’ Faces minus Places | 213 | 0.32 | 0.62 | 0.04 |
| Ce, ‘Threat-Related’ Faces minus Places | 213 | 0.27 | 0.49 | 0.03 |

^a^ Relative to the implicit baseline (i.e., certain-safety anticipation). Abbreviations—BST, bed nucleus of the stria terminalis; Ce, dorsal amygdala in the region of the central nucleus; EAc, central extended amygdala; *M*, mean; *N*, sample size; *SD*, standard deviation; *SEM*, standard error of the mean; ROI, region of interest.
