## Extended Data Table 7-2 for "Neuroticism/negative emotionality is associated with increased reactivity to uncertain threat in the bed nucleus of the stria terminalis, not the amygdala" (from another version of this preprint: version 5, 2024-04-21)

**Extended Data Table 7-2.** Inferential statistics for EAc reactivity to the threat-anticipation and threat-perception paradigms (anatomically defined ROIs; spatially unsmoothed data).

| **Contrast** | ***One-Sample Student’s t*** | ***df*** | **Nominal *p*** | **Cohen's *d*** |
| --- | --- | --- | --- | --- |
| BST, Threat minus Safety Anticipation | 9.37 | 219 | < 0.001 | 0.63 |
| Ce, Threat minus Safety Anticipation | 8.10 | 219 | < 0.001 | 0.55 |
| BST, Uncertain-Threat Anticipation^a^ | 5.50 | 219 | < 0.001 | 0.37 |
| Ce, Uncertain-Threat Anticipation^a^ | 3.10 | 219 | 0.002 | 0.21 |
| BST, Certain-Threat Anticipation^a^ | 5.44 | 219 | < 0.001 | 0.37 |
| Ce, Certain-Threat Anticipation^a^ | 3.64 | 219 | < 0.001 | 0.25 |
| BST, ‘Threat-Related’ Faces minus Places | 7.48 | 212 | < 0.001 | 0.51 |
| Ce, ‘Threat-Related’ Faces minus Places | 8.00 | 212 | < 0.001 | 0.55 |

^a^ Relative to the implicit baseline (i.e., Certain Safety anticipation). Note: The precise functional architecture of the EAc remains contentious, with some theorists suggesting that the Ce is differentially sensitive to cues and contexts associated with certain threat (‘fear’), whereas the BST is differentially sensitive to uncertain threat (‘anxiety’). To explore this hypothetical ‘double dissociation,’ we computed a standard 2 (*Region:* BST, Ce) × 2 (*Certainty:* Certain, Uncertain) repeated-measures GLM. As shown in **Figure 7** in the main report, results indicated that the BST was more strongly recruited by threat anticipation, relative to the Ce (*F*(1,219)=9.36, *p*=0.002, *d*=0.21, *BF_10_*=6.97). Neither the main effect of Certainty nor the critical Region × Certainty (‘double-dissociation’) interaction was significant (*Certainty*: *F*(1,219)=0.19, *p*=0.66, *d*=0.03, *BF_10_*=0.08; *Region × Certainty*: *F*(1,219)=0.85, *p*=0.36, *d*=0.06, *BF_10_*=0.11). Likewise, the pairwise contrast of the two threat-anticipation conditions was not significant in either region (*Ce*: *t*(219)=0.36, *p*=0.72, *d*=0.02, *BF_10_*=0.08; *BST*: *t*(219)=-0.82, *p*=0.41, *d*=-0.06, *BF_10_*=0.19). For the Ce, 48.6% participants evinced greater activation during the certain-threat anticipation (*Z_Sign_*=0.34, *p*=0.74). For the BST, 54.5% participants evinced greater activation during uncertain-threat anticipation (*Z_Sign_*=1.28, *p*=0.20). Formal tests of statistical equivalence were significant (*p*<0.001) for the omnibus Region × Certainty interaction and the focal tests of Certainty (Uncertain vs. Certain) in each region (two one-sided tests using a benchmark ‘small’ mean difference, Cohen’s *d_z_*=0.20) Collectively, these observations indicate the Ce and the BST are similarly recruited during the anticipation of temporally certain and uncertain threat, consistent with prior work by our group. Note: The data necessary to reproduce these analyses is freely available at the Open Science Framework (<https://osf.io/w5cdk>). Abbreviations—BST, bed nucleus of the stria terminalis; Ce, dorsal amygdala in the region of the central nucleus; EAc, central extended amygdala; GLM, general linear model; ROI, region of interest.
